# Evolutionary history of mammalian UDP-glucuronosyltransferase (UGT)1 and UGT2 families: the emergence of UGT2B subfamily in eutherians after the diversification of flowering plants

**DOI:** 10.1101/2021.08.20.457063

**Authors:** Yusuke K. Kawai, Kasumi Sano, Yoshinori Ikenaka, Shouta M.M. Nakayama, Mitsuki Kondo, Akira Kubota, Mayumi Ishizuka

## Abstract

The UDP-glucuronosyltransferase (UGT) gene family is responsible for the transfer of glucuronic acid to exogenous and endogenous chemicals. Based on the highly diversified number of genes, the mammalian UGT1A and UGT2B subfamily genes are believed to be involved in the conjugation reactions of xenobiotic metabolism. However, it is speculated that the UGT2 family genes are not involved in the xenobiotic metabolism of avian species due to the less diverse number of genes. In this study, we aimed to investigate the evolutionary history of mammalian UGT1 and UGT2 family genes and determine when the diversification of *UGT2B* genes occurred. We also attempted to identify the main factors responsible for the diversification of *UGT* genes and the effect of the selection pressure on the structure of the UGT isozymes. By examining the genomic information and feeding habits of 67 species representing each mammalian family, we discovered that the *UGT2B* genes emerged in the Eutheria on or after Cretaceous period and that their number were higher in plant-eating mammals (herbivore or omnivore) than in carnivorous mammals. We also found that the *UGT2B* genes in some herbivorous mammals underwent positive selection. In contrast, the diversity of the UGT1 family genes was inherited from the common ancestor of birds and mammals. Furthermore, by predicting 3D structure of UGT enzymes, estimating selection pressure on amino acid sites, and performing molecular dynamics simulations, we showed that UGT2B and some UGT2A isozymes, which have increasing gene numbers in each mammalian species, have in common that a portion of the α-helix loosens to form a hinge-like structure, that the amino acid site at which the α-helix loosens is under positive selection, and that the α-helix loosening increases the fluctuations of the UGT2B proteins. Thus, our findings suggest that the emergence of angiosperms (flowering plants) and the occurrence of “animal–plant warfare” influenced the evolution of this gene family involved in the xenobiotic metabolism of eutherians. Furthermore, future research investigating the marsupials and birds that do not possess *UGT2B* genes is required to elucidate the mechanisms underlying the metabolism of chemical substances in these species.

## Introduction

In vertebrates, the uridine diphosphate (UDP)-glycosyltransferase (UGT) superfamily conjugate sugars from UDP-sugar donors to endogenous and exogenous chemicals. Mammalian species possess the UGT1, UGT2, UGT3, and UGT8 families [1]. The UGT1 and UGT2 families, which mainly use UDP-glucuronic acid for conjugation, are called glucuronosyltransferases. The UGT1 family is composed of the UGT1A subfamily, while the UGT2 family is divided into two subfamilies, UGT2A and UGT2B [1]. Many UGT1A and UGT2B enzymes play an important role in conjugating glucuronic acids in the liver [2], while UGT2As, except for UGT2A3, are responsible for the conjugation of sugars in the olfactory neuroepithelium [3,4]. The *UGT1A* genes are composed of five exons, including the variable 1st exon and the constant 2nd–5th exons. Similarly, in humans and mice, the *UGT2A1* and *UGT2A2* genes consist of the variable 1st exons and constant 2nd–6th exons. In contrast, the *UGT2A3* and *UGT2B* genes are composed of unigenes [1].

Previous studies reported that mammalian *UGT1A* and *UGT2B* are diversified among species and may have evolved due to the influence of feeding habits. In the order Carnivora, *UGT1A6*, which plays an important role in metabolizing phenolic compounds, became a pseudogene in several pure meat-eating species [5]. Furthermore, the number of *UGT1A* genes is generally fewer in pinniped mammals than in omnivorous species [6], while the number of *UGT2B* genes is usually lower in carnivores than in omnivores [7]. It was also found that the conjugation activities of liver microsomes in carnivorous species are less than those in omnivorous species [6,7].

The diversification of *UGT2B* genes is believed to be a mammal-specific event. Similar to mammals, the number of *UGT1E* genes that are equivalent to the mammalian UGT1A genes and the conjugation activity are low in carnivorous birds. However, unlike in mammals, the number of UGT2 family genes (n ≤ 3) is generally consistent in many bird species [8,9]. These studies suggest that the mammalian and avian UGT1 and UGT2 family genes are independently diversified and evolved. In particular, the differences between mammalian UGT2B and avian UGT2 family genes imply that the ancestor *UGT2* genes did not play a role in xenobiotic metabolism and that the mammalian *UGT2B* genes gained this novel function after the divergence of mammals and birds. However, the factors that influenced the diversification of mammalian UGT1 and UGT2 family genes and the time period that this occurred are still unclear.

Previous studies have suggested that the number of *UGT* genes is related to animal feeding habits and that “animal–plant warfare” may have caused the evolution of *UGT* genes [10]. Hence, we focused on feeding habits in this study. Furthermore, herbivorous and omnivorous animals are exposed to various plants chemicals, and diets are a major source of xenobiotics. It was predicted that each plant species synthesizes 4.7 (=98,800/20,741) unique metabolites on average [11] and that ∼374,000 plant species exist on Earth [12]. Angiospermae, the class of flowering plants, is the major plant group on Earth that consists of ∼300,000 species [12]. Many herbivorous and omnivorous species consume flowering plants, which are believed to have emerged and diverged around the Upper Triassic and Cretaceous periods [13]. These suggest that the variety of chemicals to which herbivorous and omnivorous animals were exposed increased after the Cretaceous, as flowering plants diversified.

In the present study, we aimed to elucidate the evolutionary history of mammalian UGT1 and UGT2 family genes and verify the hypothesis that the development of flowering plants in the Cretaceous period caused the emergence and divergence of mammalian *UGT2B* genes. We determined the number of UGT1 and UGT2 family genes in mammals based on a public genome database by tracing their evolutionary history, investigating the relationship between the number of genes and feeding habit, and estimating selection pressures. We also investigated the properties of mammalian UGT2B by predicting its 3-dimensional (3D) structure, estimating the amino acid sites under positive selection, and investigating how it behaves in molecular dynamics simulations.

## 2. Materials and Methods

### 2.1 Homology search and genomic localization of *UGT* sequences

To determine the number of mammalian UGT1 and UGT2 family genes, we retrieved the *UGT1* and *UGT2* sequences from 67 species (Table S1) using homology searches. These species represented the 67 mammalian families with assembly information in the NCBI genome project (as of January 2020) [14] and were selected based on assembly quality, such as scaffold N50. For the homology searches, we used query sequences from the UGT1 and UGT2 family genes of seven vertebrates, namely anole lizard (*Anolis carolinensis*), Tropical clawed frog (*Xenopus tropicalis*), zebrafish (*Danio rerio*), chicken (*Gallus gallus*), platypus (*Ornithorhynchus anatinus*), mouse (*Mus muscles*), and human (*Homo sapiens*), that were annotated in Ensembl (release 98) [15] (Data S1). The TBLASTN searches [16,17] were performed for each species using the GenBank RefSeq database [18], with e-value < 1e-6 as the identity threshold. Following BLAST search, we excluded the genes with distinctly different annotations and retrieved the locus information of the remaining genes from GenBank using reutils package [19] Data visualization was performed using genoPlotR [20] from R version 4.02 [21].

### 2.2 Phylogenetic analysis of mammalian *UGT* genes

The gene location and BLASTN searches were used to classify *UGT* genes into the UGT1 and UGT2 families. Maximum likelihood-based phylogenetic analysis was performed for the classified *UGT1* and *UGT2* genes, which were divided into the variable 1st exons and other constant exons and separately analyzed for phylogeny construction. Briefly, the amino acid sequences were aligned with MAFFT version 7.2 [22] using the auto option and then trimmed with trimAl using the automated1 option [23]. For the model selection and phylogenetic analysis of *UGT1A*, the 1st and 2nd–5th exons with >200 amino acid sequences were used. For *UGT2*, the 1st and 2nd–6th exons with >200 and >250 amino acid sequences, respectively, were analyzed (Data S2). The best-fit model was selected using the Bayes information criterion (BIC) calculated by CodeML on Aminosan [24,25]. Based on the model, the phylogenies of UGT1 and UGT2 families were inferred by maximum likelihood method, with 100 bootstrap replicates, using RAxML version 8.2.10 [26]. The models used in this analysis were shown in Table S2.

### 2.3 Number of *UGT* genes

The number of UGT genes were based on the phylogenetic analysis and gene localization on the chromosome. In this study, the genes were categorized into three types: pseudogenes, partial genes, and functional genes. First, we identified the pseudogenes according to the GenBank annotation. Second, the short sequences that were excluded from the phylogenetic analysis were considered as partial genes. Third, the remaining genes were predicted to be functional genes. Fourth, with manual curation sequences containing multiple identities in one annotation were considered as multiple genes (Data S3). The mammalian phylogenetic tree was obtained from TimeTree database [27–29]. Data visualization incorporating the mammalian phylogenetic tree was performed using ggtree [30–32] from R version 4.02 [21].

### 2.4 Relationship between the number of *UGT* genes and feeding habit

The dietary information was obtained from PHYLACINE 1.2.1 [33]. The feeding habits were divided into the carnivore group that does not eat plants (0%) and the herbivore or omnivore group that eat plants (10∼100%) (Table S1). For phylogenetic correction, mammalian phylogenetic tree in TimeTree [27–29] was used and Phylogenetic ANOVA was performed to compare the number of genes between the two groups using phythools [34] from R version 4.02 [21].

### 2.5 Estimating selection pressures

The RELAX program [35] in HyPhy [36] was used to detect relaxed selection on the 1st exons of *UGT1A6* genes in carnivorous species. In the phylogenetic tree, branches corresponding to carnivorous species were considered as "test" branches, branches corresponding to herbivorous and omnivorous species were considered as "reference" branches, and ancestral branches whose diet could not be inferred were considered as "unclassified" branches to estimate the loosening of selection pressure.

The adaptive branch-site random effects likelihood (aBSREL) approach [37] in HyPhy was used to detect episodic positive selection on all branches of the 1st exon regions that encodes aglycone binding sites of eutherian or placental mammal (Infraclass Eutheria) *UGT2B* genes or marsupial (Infraclass Marsupialia/Metatheria) *UGT2* genes in herbivorous or omnivorous mammals. Positive selection was tested for branches corresponding to herbivorous and omnivorous species and for branches that were presumed to be herbivorous or omnivorous in the ancestral species. The false discovery rate (FDR) method [38] was employed for the correction of the *p-*values from all tests of branches after multiple testing. Sequences and phylogeny used in the analysis were shown in Data S4–S7.

The Fast, Unconstrained Bayesian AppRoximation (FUBAR) program [39] was employed to detect positive selection sites in 1st exon regions of *UGT1A*s and *UGT2B*s. The sequences used in the analysis were shown in Data S8 and S9 for *UGT1A*s and *UGT2B*s, respectively. The RELAX, aBSREL, and FUBAR analyses were performed in the Datamonkey server [40].

### 2.6 Predicting structure of UGT enzyme

The 3D structures of human UGTs were retrieved from AlphaFold Protein Structure Database [41]. The 3D structures of UGT enzymes of elephant (*Loxodonta africana*), armadillo (*Dasypus novemcinctus*), opossum (*Monodelphis domestica*), and platypus, which are representative species of Afrotheria, Xenarthra, Marsupialia, and Monotremes, as well as human UGT2A1 and human UGT2B4 excluding the membrane binding region and signal peptide region were predicted with AlphaFold2 [42] via ColabFold [43]. Predicted 3D structures were verified using ERRAT [44]. The protein structures were visualized with UCFC Chimera [45].

### 2.7 Molecular dynamics of human UGT2A1 and UGT2B4

Molecular dynamics (MD) simulation of human UGT2A1 and UGT2B4 without membrane binding region and signal peptide region were performed with OpenMM [46]. The protein structure after pre-processing and the configuration script used for the simulation are shown in Data S10 and S11 for UGT2A1 and UGT2B4, respectively. Root-mean-square fluctuation (RMSF), which represents the deviation at a reference position over time and is a measure of the variability of the carbon backbone (Cα atoms selected) of each UGT, was calculated using MDAnalysis [47,48].

## Results

### 3.1 Phylogeny and gene locus of UGT1 and UGT2 family genes

The maximum likelihood-based phylogenetic analysis of the variable 1st exons revealed that the *UGT1A* genes of eutherians and marsupials grouped into 4 major clades, specifically “*UGT1A1*,” “*UGT1A2*–*UGT1A5*,” “*UGT1A6*,” and “*UGT1A7*–*UGT1A10*.” Four platypus *UGT1A* genes composed one clade with the mammalian “*UGT1A1*” and “*UGT1A2*–*UGT1A5*” genes, while the remaining platypus *UGT1A* gene clustered with the avian *UGT1E* group V gene (Fig. 1, Fig. S1). Phylogenetic analysis of the constant 2nd–5th exons showed that the eutherian *UGT1A* genes formed one clade, and marsupial and platypus *UGT1A* genes composed one clade with avian *UGT1E* genes (Fig. S2). The locus information of the *UGT1A* genes showed conserved synteny among mammalian species, of which most were located between ubiquitin-specific protease 40 (*USP40*) and Transient receptor potential melastatin 8 (*TRPM8*) (Fig. 2, Fig. S3). In most species, *UGT1A* genes are usually composed of one set of 2nd–5th exons and variable 1st exons. In some species, showed duplications and deletions in the second-fifth exons, probably due to incomplete genomic information, such as in the Ord’s kangaroo rat (*Dipodomys ordii*), naked mole-rat (Heterocephalus glaber), and Chinese tree shrew (*Tupaia chinensis*) that may have multiple sets of 2nd–5th exons (Fig. S2, Fig. S3). Furthermore, we could not find the 2nd–5th *UGT1A* exons in the big brown bat (*Eptesicus fuscus*), Sunda pangolin (*Manis javanica*), northern greater galago (*Otolemur garnettii*), and black flying fox (*Pteropus alecto*).

**Figure 1.**
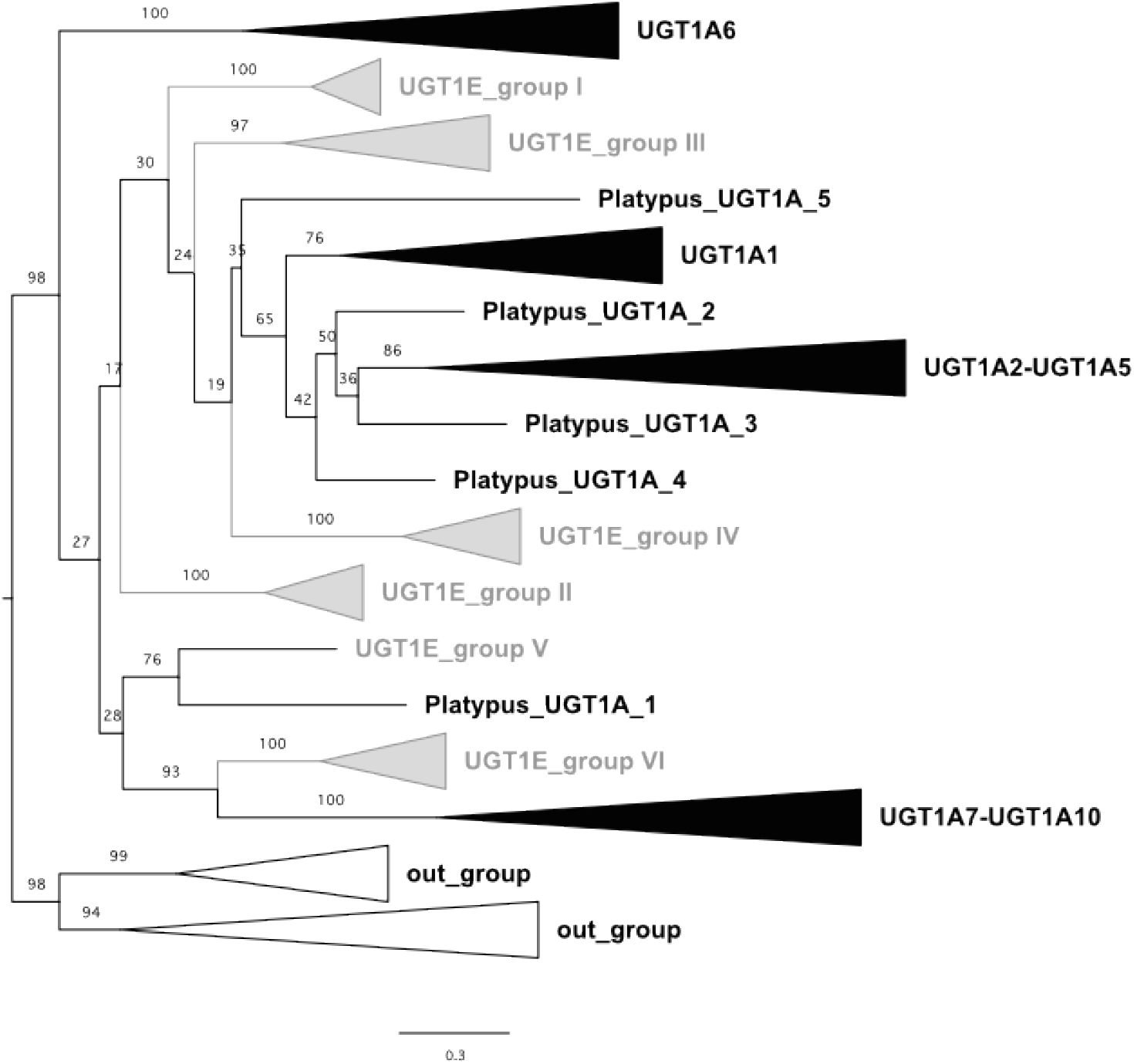
Phylogenetic tree of UDP-glucuronosyltransferase 1 (UGT1) family genes constructed from the amino acid sequence alignments of the 1st exons in mammalian and avian *UGT1* genes. The black and gray triangles indicate the mammalian *UGT1A* and avian *UGT1E* clades, respectively. The white triangles represent the outgroups. The *UGT1E* clade names were based on Kawai et al. (2018). Bootstrap values with 100 replicates are shown next to the branches as percentage. The tree is drawn to scale with branch length indicating the expected number of substitutions per site.

**Figure 2.**
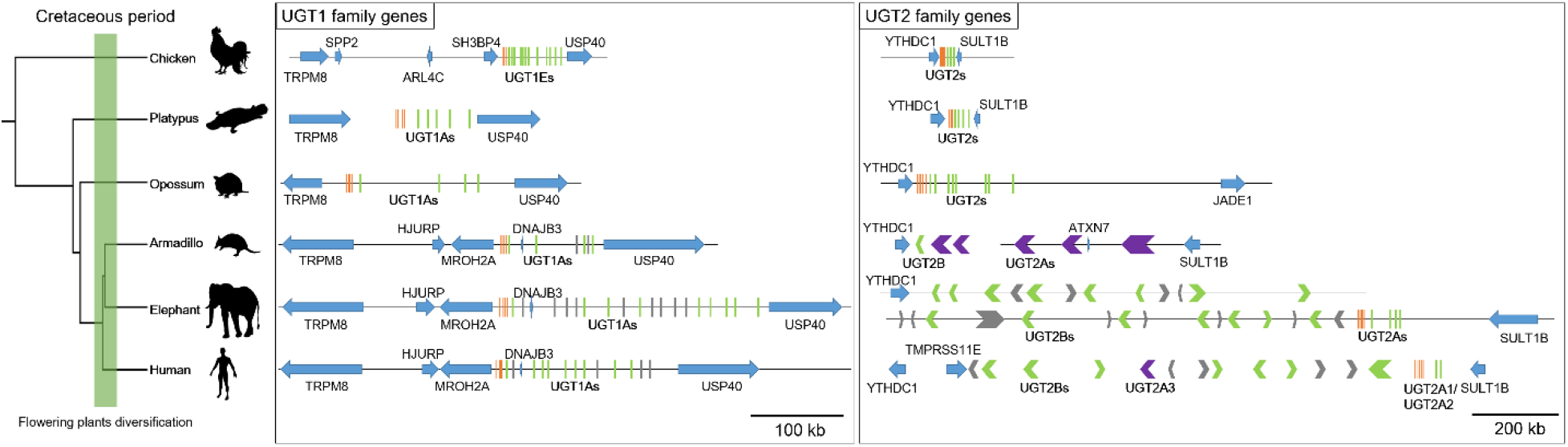
Phylogenetic tree demonstrating the synteny of UGT1 and UGT2 family genes in chicken (*Gallus gallus*), platypus (*Ornithorhynchus anatinus*), opossum (*Monodelphis domestica*), armadillo (*Dasypus novemcinctus*), elephant (*Loxodonta africana*), and human (*Homo sapiens*). *UGT* genes with multiple variable 1st exons and constant 2nd–6th exons are represented by green and orange vertical lines, respectively. The *UGT2B* and *UGT2A* unigenes are represented by green and purple arrowheads, while the other genes located in the same chromosome are represented by blue arrows. The scale bar represents 200 kilobase pairs of nucleotides.

Phylogenetic analysis of the constant 2nd–6th exons revealed that the mammalian *UGT2* genes clustered into four major clades, platypus *UGT2*, marsupial *UGT2*, eutherian *UGT2A*, and eutherian *UGT2B* (Fig. 3, Fig. S4). Moreover, the eutherian *UGT2B* clade included two subclades; *UGT2B-I* also known as *UGT2B* subfamily and *UGT2B-II* also known as *UGT2C* or *UGT2E* subfamily. The phylogenetic tree showed that the UGT2B subfamily is a unique eutherian gene group and that the 1st exons of marsupial *UGT2* genes grouped with the eutherian clades for *UGT2A* and *UGT2B-I* genes (Fig. S5). The locus information indicated the conserved synteny of *UGT2* genes among mammalian species, of which most were located between sulfotransferase family 1B member 1 (*SULT1B1*) and YTH domain-containing 1 (*YTHDC1*) (Fig. 2, Fig. S6). Similar to the *UGT1A* in all species, the marsupial *UGT2* and eutherian *UGT2A* genes were composed of variable 1st exons and constant 2nd–6th exons, while eutherian *UGT2B* genes were found to be variable unigenes.

**Figure 3.**
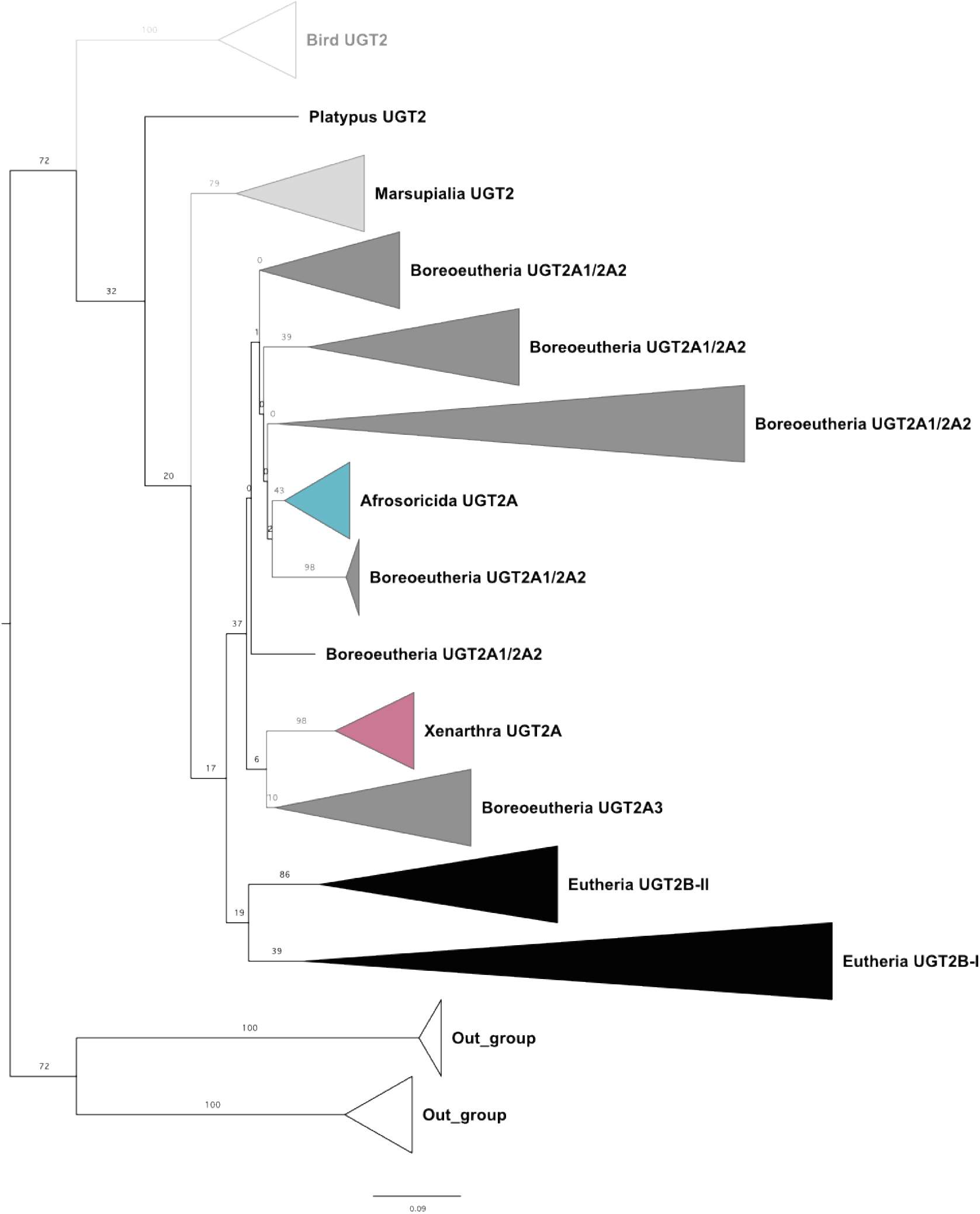
Phylogenetic tree of UDP-glucuronosyltransferase 2 (UGT2) family genes constructed from the amino acid sequence alignments of the 2nd–6th exons in mammalian and avian *UGT2* genes. The white, light gray, dark gray, blue, red, and black triangles indicate the avian (Class Aves) *UGT2*, marsupial (Infraclass Marsupialia) *UGT2*, boreoeutherian (Magnorder Boreoeutheria) *UGT2A*, afrosoricidan (Order Afrosoricida) *UGT2A*, xenarthran (Magnorder Xenarthra) *UGT2A*, and eutherian (Infraclass Eutheria) *UGT2B* clades, respectively. Bootstrap values with 100 replicates are shown next to the branches as percentage. The tree is drawn to scale with branch length indicating the expected number of substitutions per site.

### 3.2 Number of *UGT* genes

The different numbers of *UGT1A* genes in the studied mammalian species suggest that the number of functional genes varies among species, ranging from one in the minke whale (*Balaenoptera acutorostrata*) to 14 in the Common degu (*Octodon degus*) (Fig. 4). In the eutherian and marsupial *UGT1A* subgroups, the number of functional genes in “*UGT1A1*” and “*UGT1A6*” are relatively consistent, while those in “*UGT1A2*–*UGT1A5*” and “*UGT1A7*– *UGT1A10*” are variable (Fig. 5). Furthermore, the number of UGT2 family genes in mammalian species indicates the high variability of functional *UGT2* genes among mammals, which ranged from zero in the Pacific white-sided dolphin (*Lagenorhynchus obliquidens*) to 24 in the European rabbit (*Oryctolagus cuniculus*) (Fig. 4). In Eutheria, the number of *UGT2A* genes was relatively consistent compared to that of *UGT2B* genes (Fig. 6).

**Figure 4.**
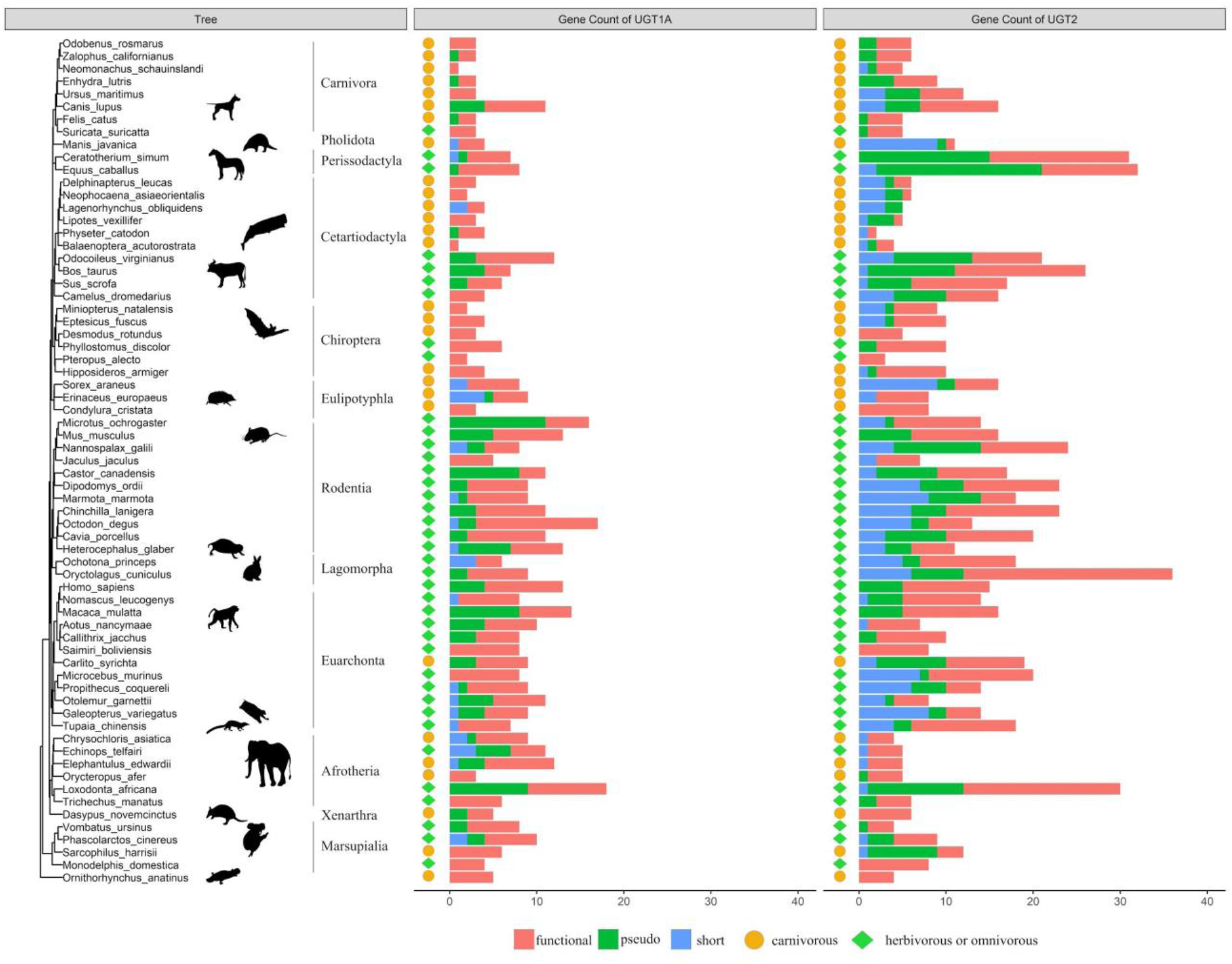
The numbers of UGT1 and UGT2 family genes among mammalian species. The data were based on GenBank annotations. The functional, pseudo-, and short genes (excluded from phylogenetic analysis) are represented by red, green, and blue bars, respectively. The feeding habit is marked on the left of the bar graphs, with orange circles and light green diamonds representing the carnivorous and herbivorous or omnivorous animals, respectively.

**Figure 5.**
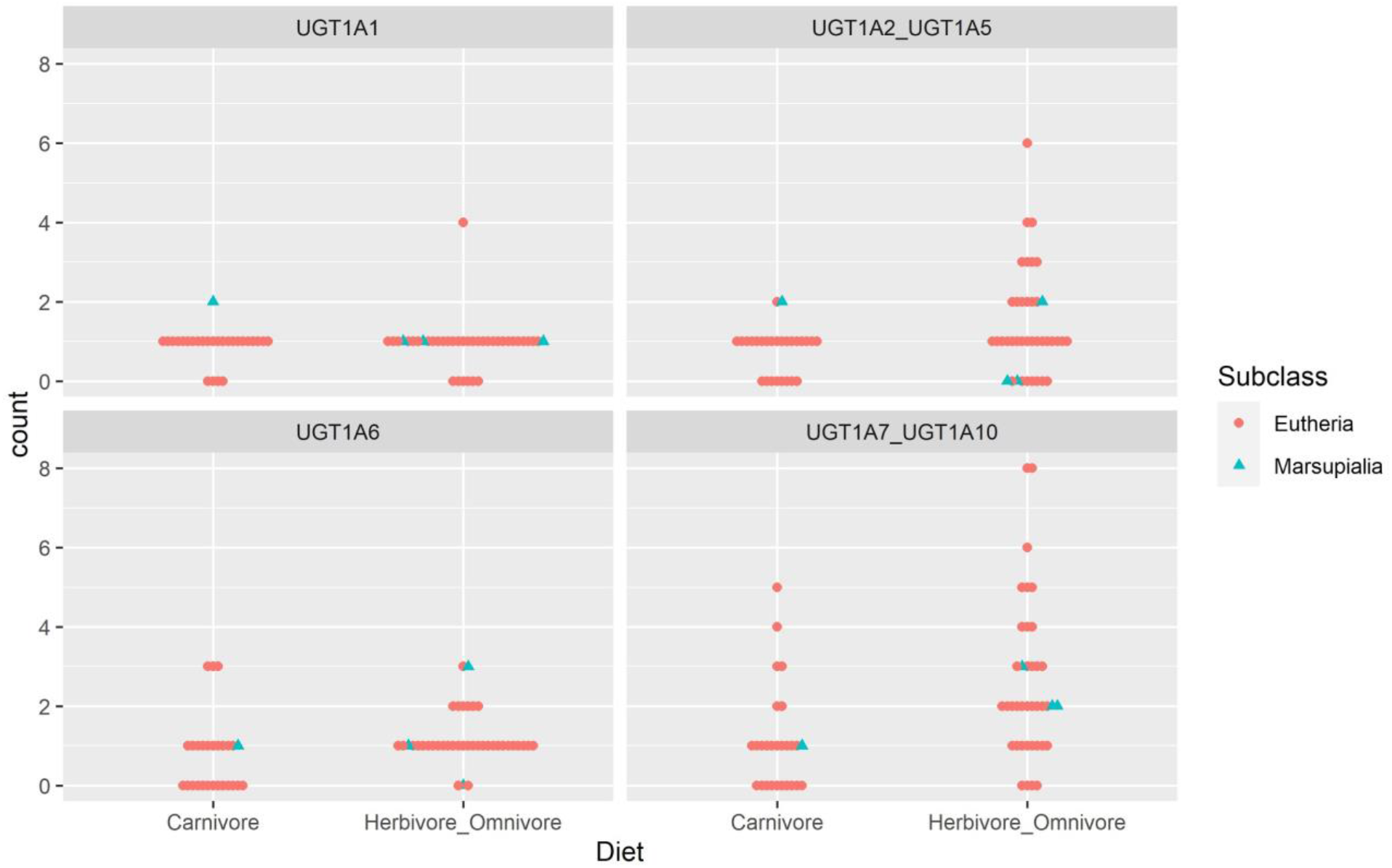
The relationship between the number of *UGT1A* genes and feeding habit. The horizontal axis indicates feeding habits and the vertical axis indicates the number of *UGT1A* genes per species, where each point represents one species. Eutherian species are indicated by orange circles and marsupial species by blue triangles. The numbers of “*UGT1A1*” and “*UGT1A6*” subgroup genes are relatively consistent, while those of “*UGT1A2*–*UGT1A5*” and “*UGT1A7*–*UGT1A10*” subgroup genes are relatively variable. There is no significant difference between the numbers of “*UGT1A*” subgroup genes in carnivorous and herbivorous or omnivorous mammals; all *P*-values are more than 0.05.

**Figure 6.**
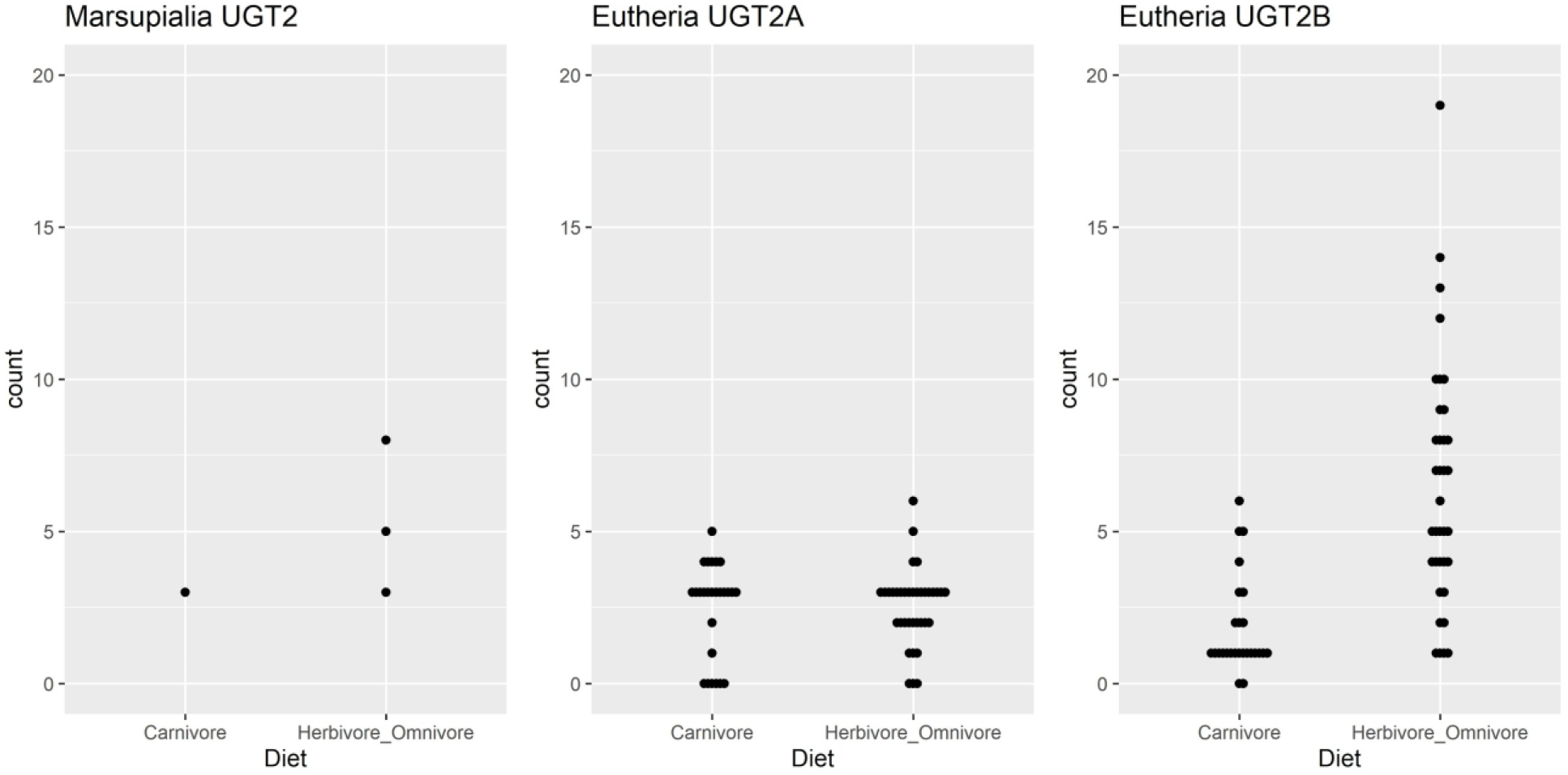
The relationship between the number of UGT2 family genes and feeding habit. The horizontal axis indicates feeding habits and the vertical axis indicates the number of *UGT2* genes per species, where each point represents one species. In Marsupialia, the number of *UGT2* genes was higher in the herbivore or omnivore group than in the carnivore group, although the sample size was small. In Eutheria, the number of *UGT2A* genes showed no significant difference between the two groups (*P* = 0.922), while the number of *UGT2B* genes was significantly higher in herbivores or omnivores than in carnivores (*P* = 0.001).

### 3.3 Relationship between the number of *UGT* genes and feeding habit

Results of the phylogenetically corrected ANOVA revealed that the number of genes in each *UGT1A* subgroup was not significantly different between carnivores and herbivores or omnivores. The analysis also indicated that the number of eutherian *UGT2A* genes were not significantly different between feeding habits. In contrast, the number of eutherian *UGT2B* genes was significantly different between feeding habits. We could not analyze the number of *UGT1A* and *UGT2* genes in Marsupialia because of the limited genomic and dietary information on marsupials. However, we observed that the number of *UGT2* genes in Tasmanian devil (*Sarcophilus harrisii*), pure carnivores, is less than or equal to that of other herbivorous or omnivorous Marsupialia species (Fig. 6). On the other hand, the number of *UGT1A* genes in Tasmanian devil show the equal to or more than that of other Marsupialia species (Fig. 5).

### 3.4 Relationship between selection pressures on *UGT* genes and feeding habit

The selection rate analysis revealed that there was no significant relaxed selection on *UGT1A6* in carnivorous mammals. In eutherian *UGT2B* genes, we found evidence of episodic diversifying selection on five out of 368 branches that were selected from the herbivorous or omnivorous mammals. The five branches consisted of the 1st exons in the *UGT2B* genes of four species, namely the pale spear-nosed bat (*Phyllostomus discolor*), European rabbit, guinea pig (*Cavia porcellus*), and Sunda flying lemur (*Galeopterus variegatus*) (Data S12, Fig. S7). In marsupial *UGT2* genes, we also found evidence of episodic diversifying selection on four out of 35 branches that were selected from the herbivorous or omnivorous mammals. All four branches included the *UGT2* genes of koala (*Phascolarctos cinereus*) (Data S13, Fig. S8).

### 3.5 Structures of UGT enzymes and estimated positive selection sites

The 3-dimensional structures of UGT enzymes were predicted with AlphaFold2 via ColabFold. The ERRAT validation showed that the overall quality factor for all predicted models was higher than 83%, while the ideal quality factor was more than 95%. In this study, an ERRAT value of higher than 90% was considered acceptable. In the predicted UGT structure model, almost all of UGT2Bs and some of UGT2As were estimated to loosen part of the helix structure to form a hinge structure (Fig. 7, Table S3). Aggregation of the predicted conformations indicate that in the predicted UGT2 family, there are at most three UGTs that maintain the helix structure in any species, whereas there are many UGTs that have a hinge structure.

**Figure 7.**
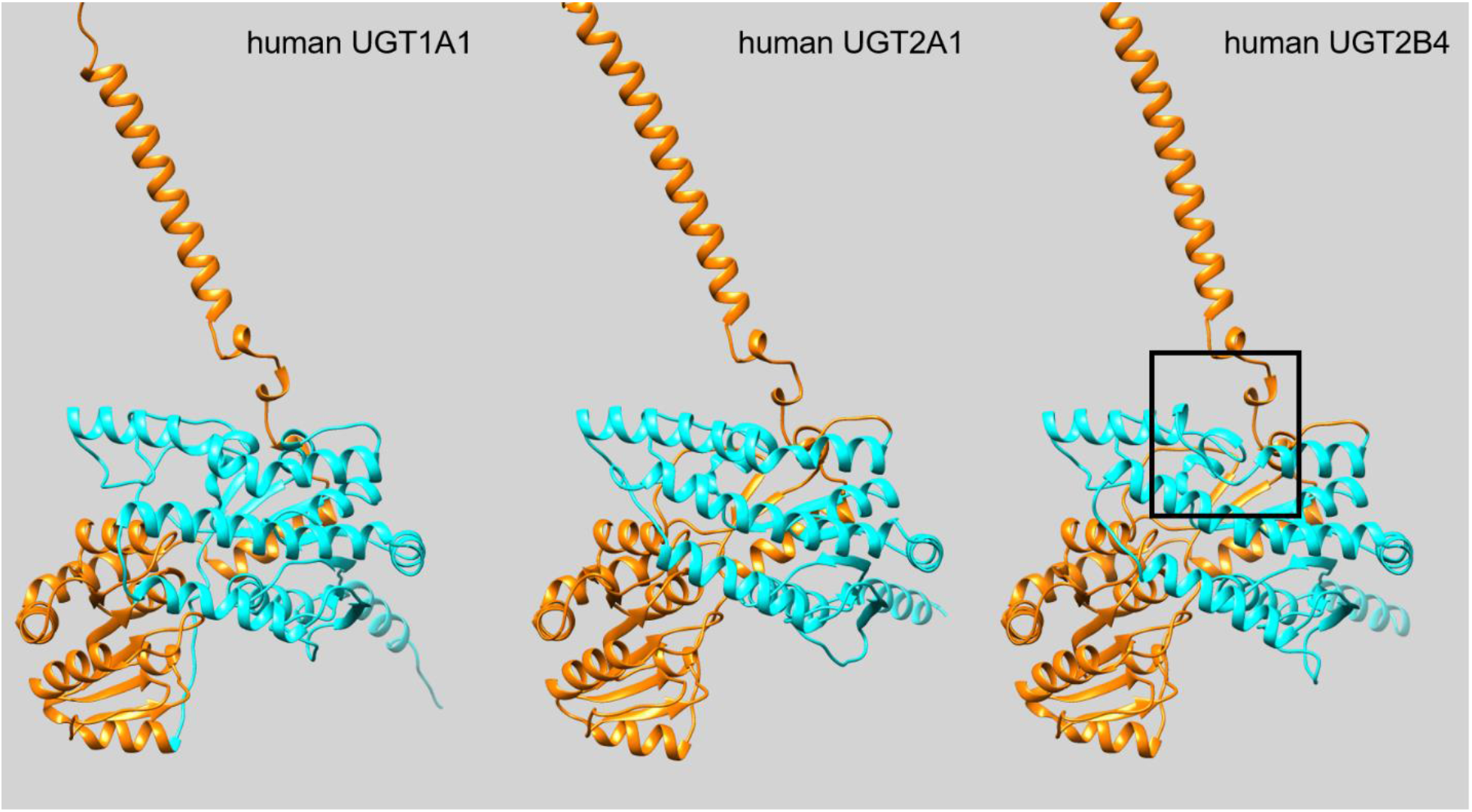
Predicted UGT structure with AlphaFold2. Representative structures of human UGT1A1 (UniProt accession: Q5DT03), UGT2A1 (UniProt accession: P0DTE4), and UGT2B4 (UniProt accession: P06133) are shown. The N-terminal part encoded by the first exon is shown in light blue, and the C-terminal part encoded by the second exon and below is shown in orange. UGT2B4 has a hinge structure with a loosened helix structure in the box region, and this hinge structure is observed in almost all UGT2B and some UGT2A.

In addition, the predicted models were used to confirm the conformational position of the amino acid residues that are predicted to be under positive selection pressure in each family. FUBAR analysis in Datamonkey server estimated 5 sites undergoing positive selection with posterior probability is over 0.9 within the region of 1st exon of the UGT1A subfamily genes. The 5 estimated positive selection sites corresponded to Asn 98, Asn 102, Ile 116, Val 225, and Gln 288 in human UGT1A1, respectively (Fig. 8a). FUBAR analysis also found 10 sites undergoing positive selection with posterior probability is over 0.9 within the region of 1st exon of the UGT2B subfamily genes. The 10 estimated positive selection sites corresponded to Leu 81, Phe 86, Leu 93, Arg 96, Gln 111, Thr 118, Phe 119, Arg 124, Val 153, and Ile 226 in human UGT2B4, respectively (Fig. 8b). In particular, the region where the helix structure loosens in UGT2Bs was a positive selection target (Ile 226).

**Figure 8.**
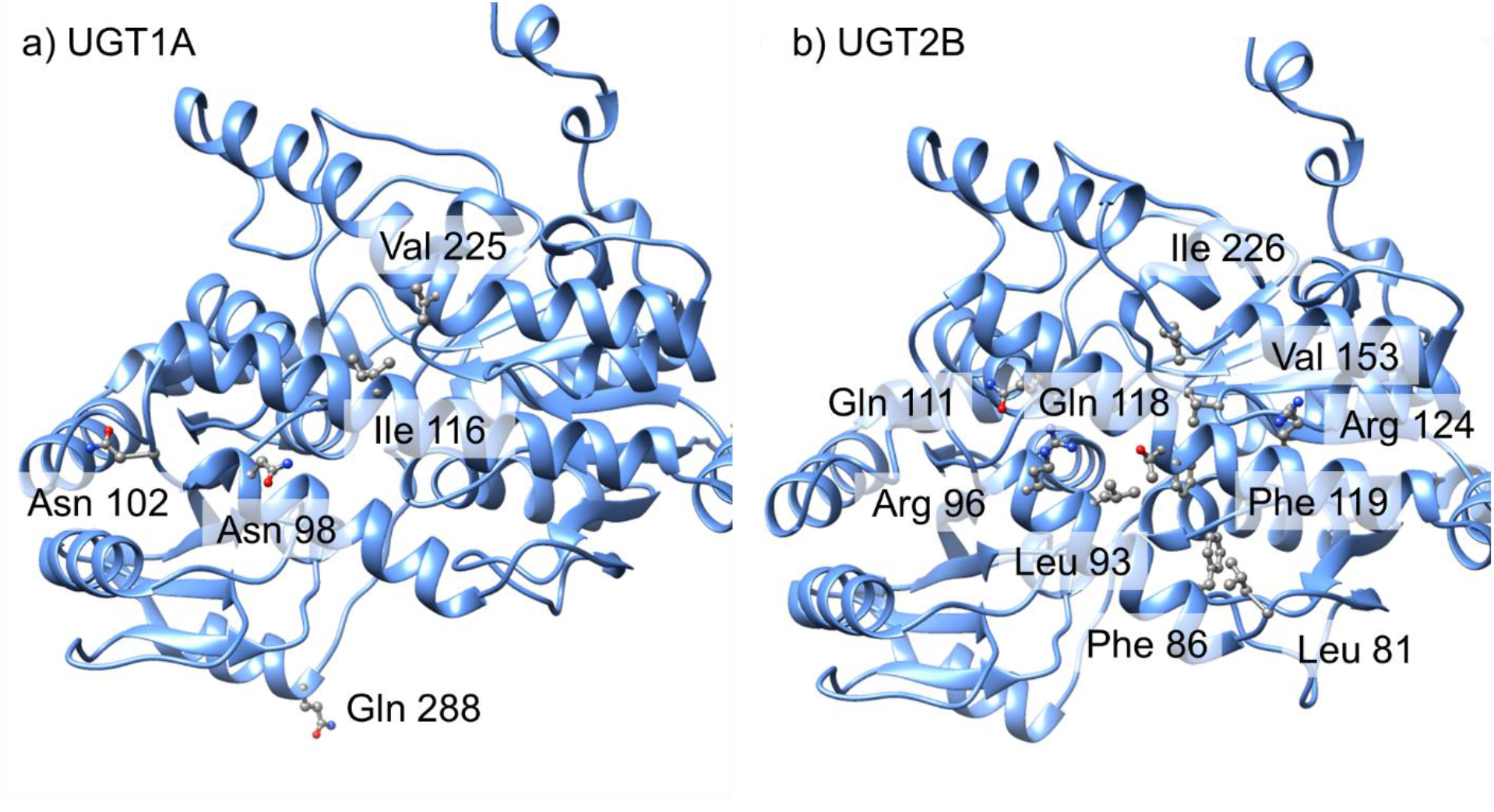
Positive selection sites in the N-terminal domain of UGT estimated by FUBAR analysis. a) Five estimated positive selection sites in mammalian UGT1As are shown on the structure of human UGT1A1. b) Ten estimated positive selection sites in mammalian UGT2Bs are shown on the structure of human UGT2B4. The region where the helix structure loosens in UGT2Bs was a positive selection target (Ile 226).

Moreover, to estimate how the loosening of the center of the helix region affected the behavior of UGT, MD simulations using OpenMM were performed based on predicted models of UGT2A1 and UGT2B4 excluding the membrane binding region and the signal peptide region. The calculated RMSF of the backbone Cα atom showed that UGT2B4 has a larger fluctuation from Thr 204 to Leu 240 in the hinge structure compared to UGT2A1, and also showed that UGT2B4 has a larger fluctuation compared to UGT2A1 in the overall structure (Fig. S9).

## Discussion

The locus information indicated that almost all *UGT1A* genes were composed of variable 1st exons and constant 2nd–5th exons. Furthermore, the evolutionary history of *UGT1A* genes was deduced from the phylogenetic analysis of the 1st exons, revealing that the common ancestor of mammals and birds possessed at least seven UGT1 family genes. This is in agreement with a previous report on the common ancestor of vertebrates possessing multiple UGT1 family genes [49]. The phylogenetic analysis also indicated that the UGT1A subfamily genes would show the birth–death evolution after the divergence of mammals and birds [50].

The *UGT1A* genes of marsupials and eutherians can be divided into four subgroups, namely “*UGT1A1*,” “*UGT1A2*–*UGT1A5*,” “*UGT1A6*,” and “*UGT1A7*–*UGT1A10*”, corresponding to the human *UGT1A* [1]. The number of *UGT1A* genes were diversified among species. However, the diversification patterns differed among the *UGT1A* subgroups. For example, the “*UGT1A1*” and “*UGT1A6*” subgroups were consistent among mammalian species. In particular, *UGT1A1* was found to be conserved in almost all mammals. In contrast, the “*UGT1A2*– *UGT1A5*” and “*UGT1A7*–*UGT1A10*” subgroups showed diversity among mammalian species. Therefore, the common ancestor of eutherians and marsupials might have possessed four UGT1A subgroups, with their functions conserved throughout mammalian evolutionary history.

Our results suggest that UGT1A1, which is known to conjugate bilirubin [51], was important for mammalian species; hence, the number of UGT1A1 was conserved during evolution. However, we did not find *UGT1A1* ortholog in the current platypus genomic information. This suggests that another *UGT1A* ortholog may be involved in the bilirubin conjugation in platypus or that this may be due to incomplete platypus genomic information.

Traditionally, it was thought that human UGT1A2–UGT1A5 enzymes conjugate endogenous substrates, such as bile acids, while UGT1A7–UGT1A10 conjugate exogenous substances, such as phenol [52,53]. However, UGT1A3, which was found in the “*UGT1A2*– *UGT1A5*” subgroup, reportedly conjugates other exogenous chemicals, such as amines, flavonoids, and hydroxylated benzo[a]pyrene [51]. Furthermore, our results suggest that the enzymes in the “*UGT1A2*–*UGT1A5*” subgroup are required for the metabolism of exogenous chemicals. However, the number of genes in the “*UGT1A2*–*UGT1A5*” subgroup was not significantly different between carnivorous and herbivorous or omnivorous mammals.

UGT1A6 is a major drug-metabolizing enzyme [54], with its gene known as a pseudogene in numerous carnivorous species [5,6]. However, UGT1A6 was not duplicated in many herbivorous or omnivorous species. In the present study, we did not observe significant differences between the numbers of genes of the *UGT1A* subgroups. There was also no relaxation of selection pressure on *UGT1A6* in carnivorous species. These results suggest that *UGT1A6* is partially involved in the metabolism of plant chemicals especially in the order of Carnivora but mainly involved in the conjugation of exogenous chemicals that existed before the emergence of flowering plants, such as benzo[a]pyrene metabolites in other eutherian orders [54].

The “*UGT1A7*–*UGT1A10*” subgroup genes were discovered to be more abundant in herbivores or omnivores than in carnivores, although there was no significant difference between the two. In humans, UGT1A7–UGT1A10 conjugate not only carcinogenic compounds [55,56] but also dietary compounds, such as flavonols and isoflavones [57,58]. Therefore, the production of new plant chemicals may have caused an increase in the number of “*UGT1A7*–*UGT1A10*” subgroup genes in herbivorous or omnivorous mammals.

The locus information revealed that some *UGT2A* genes were composed of variable 1st exons and constant 2nd–6th exons, while other *UGT2A* and *UGT2B* genes were unigenes composed of 1st–6th exons. Therefore, we deduced the evolutionary history of UGT2 family genes from the phylogenetic analysis of the constant 2nd–6th exons and discovered that mammalian *UGT2* genes grouped into four major clades, namely the platypus *UGT2*, marsupial *UGT2*, eutherian *UGT2A*, and eutherian *UGT2B*. This result suggests that after the divergence of Marsupialia and Eutheria, the UGT2 family was divided into two eutherian subgroups, UGT2A and UGT2B. Moreover, the appearance of *UGT2* genes in platypus and marsupials, which were composed of variable 1st exons and constant 2nd–6th exons, suggested that *UGT2* unigene duplication occurred during eutherian evolution.

The differences in the number of genes per clade, in which the number of UGT2B genes was higher in herbivores or omnivores than in carnivores, suggest that the diversification of eutherian UGT2B genes was influenced by feeding habits. In Eutheria, *UGT2B* genes function in the metabolism of steroid hormones in the liver [59,60] and some are known to metabolize flavonoids [61] that are known to be steroid-like chemicals [62].Therefore, this difference suggests that the increase in the number of *UGT2B* genes may have resulted from the need to metabolize exogenous chemicals in food, including flavonoids. Furthermore, marsupial *UGT2* genes may have also diversified due to changes in feeding habits. However, because of the limited number of species and incomplete genome assembly, we could not confirm its correlation to feeding habits.

The number of *UGT2A* genes was consistent among mammalian species (n ≤ 4), except in armadillo, European rabbit, and Chinese tree shrew. In humans, the *UGT2A1* and *UGT2A2* genes function during the local detoxification of polycyclic aromatic hydrocarbons (PAHs) in the olfactory neuroepithelium [3,63]. Therefore, the consistent number of eutherian *UGT2A* genes indicate that these are not related to the conjugation of food chemicals. In contrast, the human *UGT2A3*, a unigene composed of 1st–6th exons, is primarily expressed in the intestine, liver, and kidney [4,64]. Therefore, the *UGT2A* genes in armadillo, European rabbit, and Chinese tree shrew, which were duplicated similar to *UGT2B* genes, may be specifically associated with xenobiotic metabolism in the liver.

Plants synthesize many secondary metabolites for protection against herbivorous animals, including vertebrates and insects [65]. Flowering plants (Class Angiospermae) are a major group that diversified during the Cretaceous period and became a primary food source on Earth [13]. Before the Cretaceous period, Eutheria and Marsupialia diverged, and both mammalian groups were independently required to ingest and metabolize the secondary metabolites from flowering plants for continued survival.

In this study, we hypothesized that the *UGT1A* genes may have played a role in the xenobiotic metabolism in the common ancestor of mammals before the emergence of flowering plants. Although some UGT1A isozymes are known to conjugate the secondary metabolites of flowering plants, such as cannabinol and nicotine [66,67], we did not observe a relationship between the number of UGT1A subgroup genes and feeding habit. This weak relationship suggests that ancestral *UGT1A* genes might have conjugated certain chemicals, such as polycyclic aromatic hydrocarbons (PAHs), and toxic metabolites from the ferns (Class Gymnospermae) that existed before Cretaceous period and there are historical constraints in the *UGT1A* genes.

However, a previous study suggested that the *UGT2* genes in the common ancestor of mammals do not have diversity and do not play a role in hepatic xenobiotic metabolism [8]. In this study, phylogenetic analysis of the 1st exons in *UGT2* genes that is related to aglycone recognition [68], reveals that marsupial *UGT2* genes, which are composed of variable 1st exons and constant exons similar to eutherian *UGT2A1* and *UGT2A2*, formed two different clades with eutherian *UGT2A* and *UGT2B* genes. This result suggests that in the common ancestor of Eutheria and Marsupialia possessed both *UGT2A*-like and *UGT2B*-like 1st exons. Moreover, the phylogenetic tree of the 2nd–6th exons showed *UGT2A* and *UGT2B* genes were separated in the eutherian lineage. These phylogenetic inferences suggest that in the eutherian lineage 2nd– 6th exons were duplicated and separated from original *UGT2* genes with *UGT2B*-like 1st exon as unigenes. Unlike splicing variant genes such as *UGT1* family genes, unigene of *UGT2B*, which means that the distance between the 1st exon and the 2nd exon does not separate even when the number of genes increases, may have created room for more genes to be generated by duplication. The phylogenetic tree of the 2nd–6th exons also demonstrate that several duplications of *UGT2B* genes separately occurred in each eutherian family. This suggests that *UGT2B* genes have independently evolved in each lineage to facilitate the adaptation to species-specific environments and that the common ancestor of eutherian mammals “invented” the *UGT2B* genes as unigenes for metabolizing variable flowering plant chemicals.

Furthermore, in each lineage, the function of the UGT2B subgroup expanded for conjugating species-specific xenobiotics. In Eutheria, there are some exceptions to this hypothesis, such as the armadillos that possess diversified *UGT2A* genes, not *UGT2B*. Armadillos are carnivorous mammals that mainly eats small animals, such as insects [69]. Therefore, the high number of *UGT2A* genes in armadillo may have been caused by specific endogenous chemicals. Furthermore, since armadillos are xenarthran mammals that diverged from other eutherians in the Cretaceous period [70], this species might reflect the ambiguity of the functions between ancestral *UGT2B* and *UGT2A* in xenobiotic metabolism.

We tested the hypothesis that exposure to flowering plant chemicals was the major selection pressure on the 1st exons of *UGT2* genes in Eutheria and Marsupialia using aBSREL analysis. Our data suggest that there was no positive selection on the *UGT2B* genes in several herbivorous eutherian species. This may have been caused by the huge number of *UGT2B* genes analyzed, and the corrected multiple comparison tests reduced the power of detection. However, we discovered that there was episodic diversifying selection in four herbivorous or omnivorous species, which supports our hypothesis that the plant chemicals in food may have influenced the divergence of *UGT2B* genes. Moreover, we found evidence of episodic diversifying selection on the branches of *UGT2* genes in koala. Koalas are mammals that eat the leaves of eucalyptus trees that contain toxic chemicals, such as phenolic compounds and terpenes [71,72]. Although the genome assembly is incomplete, the selection evidence suggests that the *UGT2* genes in koala may have also evolved to adapt to these plant chemicals. Thus, our results suggest that convergent/parallel evolution occurred between the *UGT2* genes of marsupials and eutherians after adaptation to conjugating plant chemicals.

Based on our predicted model of the UGTs, we inferred that UGTs in which the middle part of the α-helix loosens and assumes a hinge conformation have emerged in the UGT2 family. Although the number of species for UGT structure predictions in this study was small and some of the predicted structure were unreliable, our results suggest that in the mammalian UGT2 family, the number of UGTs in which the helix structure is limited to about three in any species, while the number of UGTs with hinge structures shows much more diverse among species. In addition, we estimated the amino acid sites of positive selection in UGT2B, and found that positive selection occurred on the amino acid site where the α-helix was loosened, suggesting that the structure with a loosened α-helix, represented by UGT2B, is an important structure that is maintained despite the diversification of amino acid residues among species.

Furthermore, MD simulations of human UGT2A1 and UGT2B4 suggest that the loosening of the α-helix increases the overall fluctuation of UGT structure. These results suggest that the structure represented by UGT2B acquired more flexibility than the α-helix-maintained structure represented by UGT2A, thereby adapting to the increasing number of chemicals derived from angiosperms.

In this study we used publicly available genomic information for our analysis. One species was selected from each family, and when multiple species existed, the species with the best assembly was selected based on information such as Scaffold N50. However, the quality of the genome assembly varied, and it is possible that some of the results shown in this study are due to incomplete genome information. Some of the results due to differences in genome quality may have corrected using statistical analysis with 67 species, but further analysis using better genome information is desirable in the future. In addition, this study is an analysis based on gene sequences, and there are limitations in inferring functions from gene sequences. It is necessary to verify the evolutionary history of the UGT family suggested in this study by adding experimental data in the future.

In summary, we traced the evolutionary history of mammalian *UGT1A* and *UGT2* genes by analyzing their phylogenies, determining the relationship between number of genes and feeding habit, and estimating selection pressures. The results revealed that *UGT2B* genes emerged in Eutheria after the divergence from Marsupialia. Our findings also suggest that parallel evolution may have occurred in the *UGT2* genes of eutherian and marsupial lineages after the diversification of flowering plants in the Cretaceous period.

## Supporting information

Suppementary Datasets

## Acknowledgments

This work was supported by Grants-in-Aid for Scientific Research from the Ministry of Education, Culture, Sports, Science, and Technology of Japan (MEXT) awarded to M. Ishizuka (No. 18K19847 and 21H04919), and Hokkaido University SOUSEI Support Program. This research was also supported by Program for supporting introduction of the new sharing system (JPMXS0420100619). This research was also supported by MEXT awarded to Y. Kawai (No. 20K15848). The English in this manuscript was proofread by Editage (https://www.editage.jp/).

## References

1. Mackenzie PI, Bock KW, Burchell B, Guillemette C, Ikushiro S-I, Iyanagi T, et al. Nomenclature update for the mammalian UDP glycosyltransferase (UGT) gene superfamily. Pharmacogenet Genomics. 2005;15: 677–685.

2. Rowland A, Miners JO, Mackenzie PI. The UDP-glucuronosyltransferases: Their role in drug metabolism and detoxification. Int J Biochem Cell Biol. 2013;45: 1121–1132.

3. Jedlitschky G, Cassidy AJ, Sales M, Pratt N, Burchell B. Cloning and characterization of a novel human olfactory UDP-glucuronosyltransferase. Biochem J. 1999;340 (Pt 3): 837– 843.

4. Court MH, Hazarika S, Krishnaswamy S, Finel M, Williams JA. Novel Polymorphic Human UDP-glucuronosyltransferase 2A3: Cloning, Functional Characterization of Enzyme Variants, Comparative Tissue Expression, and Gene Induction. Mol Pharmacol. 2008;74: 744–754.

5. Shrestha B, Reed JM, Starks PT, Kaufman GE, Goldstone JV, Roelke ME, et al. Evolution of a major drug metabolizing enzyme defect in the domestic cat and other felidae: phylogenetic timing and the role of hypercarnivory. PLoS One. 2011;6: e18046.

6. Kakehi M, Ikenaka Y, Nakayama SMM, Kawai YK, Watanabe KP, Mizukawa H, et al. Uridine Diphosphate-Glucuronosyltransferase (UGT) Xenobiotic Metabolizing Activity and Genetic Evolution in Pinniped Species. Toxicol Sci. 2015;147: 360–369.

7. Kondo T, Ikenaka Y, Nakayama SMM, Kawai YK, Mizukawa H, Mitani Y, et al. Uridine Diphosphate-Glucuronosyltransferase (UGT) 2B Subfamily Interspecies Differences in Carnivores. Toxicol Sci. 2017;158: 90–100.

8. Kawai YK, Ikenaka Y, Ishizuka M, Kubota A. The evolution of UDP-glycosyl/ glucuronosyltransferase 1E (UGT1E) genes in bird lineages is linked to feeding habits but UGT2 genes is not. PLoS One. 2018;13: 1–13.

9. Kawai YK, Shinya S, Ikenaka Y, Saengtienchai A, Kondo T, Darwish WS, et al. Characterization of function and genetic feature of UDP-glucuronosyltransferase in avian species. Comparative Biochemistry and Physiology Part - C: Toxicology and Pharmacology. 2019;217: 5–14.

10. Bock KW. Vertebrate UDP-glucuronosyltransferases: functional and evolutionary aspects. Biochem Pharmacol. 2003;66: 691–696.

11. Afendi FM, Okada T, Yamazaki M, Hirai-Morita A, Nakamura Y, Nakamura K, et al. KNApSAcK family databases: integrated metabolite-plant species databases for multifaceted plant research. Plant Cell Physiol. 2012;53: e1.

12. Christenhusz MJM, Byng JW. The number of known plants species in the world and its annual increase. Phytotaxa. 2016;261: 201–217.

13. Li HT, Yi TS, Gao LM, Ma PF, Zhang T, Yang JB, et al. Origin of angiosperms and the puzzle of the Jurassic gap. Nature Plants. 2019;5: 461–470.

14. Kitts PA, Church DM, Thibaud-Nissen F, Choi J, Hem V, Sapojnikov V, et al. Assembly: a resource for assembled genomes at NCBI. Nucleic Acids Res. 2016;44: D73–80.

15. Cunningham F, Achuthan P, Akanni W, Allen J, Amode MR, Armean IM, et al. Ensembl 2019. Nucleic Acids Res. 2018;47: D745–D751.

16. Camacho C, Coulouris G, Avagyan V, Ma N, Papadopoulos J, Bealer K, et al. BLAST plus: architecture and applications. BMC Bioinformatics. 2009;10: 1.

17. Altschul SF, Gish W, Miller W, Myers EW, Lipman DJ. Basic local alignment search tool. J Mol Biol. 1990;215: 403–410.

18. O’Leary NA, Wright MW, Brister JR, Ciufo S, Haddad D, McVeigh R, et al. Reference sequence (RefSeq) database at NCBI: current status, taxonomic expansion, and functional annotation. Nucleic Acids Res. 2016;44: D733–45.

19. Schofl G. reutils: Talk to the NCBI EUtils. 2016. Available: https://CRAN.R-project.org/package=reutils

20. Guy L, Kultima JR, Andersson SGE, Quackenbush J. GenoPlotR: comparative gene and genome visualization in R. Bioinformatics. 2011;27: 2334–2335.

21. R Core Team. R: A Language and Environment for Statistical Computing. 2020. Available: https://www.R-project.org/

22. MAFFT Multiple Sequence Alignment Software Version 7 : Improvements in Performance and Usability Article Fast Track.

23. Capella-Gutiérrez S, Silla-Martínez JM, Gabaldón T. trimAl: A tool for automated alignment trimming in large-scale phylogenetic analyses. Bioinformatics. 2009;25: 1972–1973.

24. Tanabe AS. Kakusan4 and Aminosan: Two programs for comparing nonpartitioned, proportional and separate models for combined molecular phylogenetic analyses of multilocus sequence data. Mol Ecol Resour. 2011;11: 914–921.

25. Yang Z. PAML 4: phylogenetic analysis by maximum likelihood. Mol Biol Evol. 2007;24: 1586–1591.

26. Stamatakis A. RAxML version 8: a tool for phylogenetic analysis and post-analysis of large phylogenies. Bioinformatics. 2014;30: 1312–1313.

27. Hedges SB, Dudley J, Kumar S. TimeTree: a public knowledge-base of divergence times among organisms. Bioinformatics. 2006;22: 2971–2972.

28. Hedges SB, Marin J, Suleski M, Paymer M, Kumar S. Tree of life reveals clock-like speciation and diversification. Mol Biol Evol. 2015;32: 835–845.

29. Kumar S, Stecher G, Suleski M, Hedges SB. TimeTree: a resource for timelines, timetrees, and divergence times. Mol Biol Evol. 2017;34: 1812–1819.

30. Yu G, Smith DK, Zhu H, Guan Y, Lam TT-Y. Ggtree : An r package for visualization and annotation of phylogenetic trees with their covariates and other associated data. Methods Ecol Evol. 2017;8: 28–36.

31. Yu G, Lam TT-Y, Zhu H, Guan Y. Two Methods for Mapping and Visualizing Associated Data on Phylogeny Using Ggtree. Mol Biol Evol. 2018;35: 3041–3043.

32. Yu G. Using ggtree to Visualize Data on Tree-Like Structures. Curr Protoc Bioinformatics. 2020;69: e96.

33. Faurby S, Davis M, Pedersen RØ, Schowanek SD, Antonelli A, Svenning J-C. PHYLACINE 1.2: The Phylogenetic Atlas of Mammal Macroecology. Ecology. 2018;99: 2626.

34. Revell LJ. phytools: an R package for phylogenetic comparative biology (and other things): phytools: R package. Methods Ecol Evol. 2012;3: 217–223.

35. Wertheim JO, Murrell B, Smith MD, Kosakovsky Pond SL, Scheffler K. RELAX: detecting relaxed selection in a phylogenetic framework. Mol Biol Evol. 2015;32: 820–832.

36. Pond SLK, Frost SDW, Muse SV. HyPhy: hypothesis testing using phylogenies. Bioinformatics. 2004;21: 676–679.

37. Smith MD, Wertheim JO, Weaver S, Murrell B, Scheffler K, Kosakovsky Pond SL. Less is more: an adaptive branch-site random effects model for efficient detection of episodic diversifying selection. Mol Biol Evol. 2015;32: 1342–1353.

38. Controlling the False Discovery Rate : A Practical and Powerful Approach to Multiple Testing Author (s).

39. Murrell B, Moola S, Mabona A, Weighill T, Sheward D, Kosakovsky Pond SL, et al. FUBAR: a fast, unconstrained bayesian approximation for inferring selection. Mol Biol Evol. 2013;30: 1196–1205.

40. Weaver S, Shank SD, Spielman SJ, Li M, Muse SV, Kosakovsky Pond SL. Datamonkey 2.0: A Modern Web Application for Characterizing Selective and Other Evolutionary Processes. Mol Biol Evol. 2018;35: 773–777.

41. Varadi M, Anyango S, Deshpande M, Nair S, Natassia C, Yordanova G, et al. AlphaFold Protein Structure Database: massively expanding the structural coverage of protein-sequence space with high-accuracy models. Nucleic Acids Res. 2022;50: D439–D444.

42. Jumper J, Evans R, Pritzel A, Green T, Figurnov M, Ronneberger O, et al. Highly accurate protein structure prediction with AlphaFold. Nature. 2021;596: 583–589.

43. Mirdita M, Schütze K, Moriwaki Y, Heo L, Ovchinnikov S, Steinegger M. ColabFold: making protein folding accessible to all. Nat Methods. 2022;19: 679–682.

44. Colovos C, Yeates TO. Verification of protein structures: patterns of nonbonded atomic interactions. Protein Sci. 1993;2: 1511–1519.

45. Pettersen EF, Goddard TD, Huang CC, Couch GS, Greenblatt DM, Meng EC, et al. UCSF Chimera - A visualization system for exploratory research and analysis. J Comput Chem. 2004;25: 1605–1612.

46. Eastman P, Swails J, Chodera JD, McGibbon RT, Zhao Y, Beauchamp KA, et al. OpenMM 7: Rapid development of high performance algorithms for molecular dynamics. PLoS Comput Biol. 2017;13: e1005659.

47. Michaud-Agrawal N, Denning EJ, Woolf TB, Beckstein O. MDAnalysis: a toolkit for the analysis of molecular dynamics simulations. J Comput Chem. 2011;32: 2319–2327.

48. Gowers R, Linke M, Barnoud J, Reddy T, Melo M, Seyler S, et al. MDAnalysis: A python package for the rapid analysis of molecular dynamics simulations. In: Benthall S, Rostrup S, editors. Proceedings of the 15th Python in Science Conference. SciPy; 2016. doi:10.25080/majora-629e541a-00e

49. Huang H, Wu Q. Cloning and comparative analyses of the zebrafish Ugt repertoire reveal its evolutionary diversity. PLoS One. 2010;5: e9144.

50. Nei M, Rooney AP. Concerted and birth-and-death evolution of multigene families. Annu Rev Genet. 2005;39: 121–152.

51. Meech R, Hu DG, McKinnon RA, Mubarokah SN, Haines AZ, Nair PC, et al. The UDP-Glycosyltransferase (UGT) Superfamily: New Members, New Functions, and Novel Paradigms. Physiol Rev. 2019;99: 1153–1222.

52. Emi Y, Ikushiro S, Iyanagi T. Drug-responsive and tissue-specific alternative expression of multiple first exons in rat UDP-glucuronosyltransferase family 1 (UGT1) gene complex. J Biochem. 1995;117: 392–399.

53. Trottier J, Verreault M, Grepper S, Monté D, Bélanger J, Kaeding J, et al. Human UDP-glucuronosyltransferase (UGT)1A3 enzyme conjugates chenodeoxycholic acid in the liver. Hepatology. 2006;44: 1158–1170.

54. Bock KW, Köhle C. UDP-glucuronosyltransferase 1A6: structural, functional, and regulatory aspects. Methods Enzymol. 2005;400: 57–75.

55. Lacko M, Roelofs HMJ, te Morsche RHM, Voogd AC, Ophuis MBO, Peters WHM, et al. Genetic polymorphisms in the tobacco smoke carcinogens detoxifying enzyme UGT1A7 and the risk of head and neck cancer. Head Neck. 2009;31: 1274–1281.

56. Dellinger RW, Fang J-L, Chen G, Weinberg R, Lazarus P. Importance of UDP-glucuronosyltransferase 1A10 (UGT1A10) in the detoxification of polycyclic aromatic hydrocarbons: decreased glucuronidative activity of the UGT1A10139Lys isoform. Drug Metab Dispos. 2006;34: 943–949.

57. Niemeyer ED, Brodbelt JS. Regiospecificity of human UDP-glucuronosyltransferase isoforms in chalcone and flavanone glucuronidation determined by metal complexation and tandem mass spectrometry. J Nat Prod. 2013;76: 1121–1132.

58. Ruan J-Q, Yan R. Regioselective glucuronidation of the isoflavone calycosin by human liver microsomes and recombinant human UDP-glucuronosyltransferases. Chem Biol Interact. 2014;220: 231–240.

59. Locuson CW, Tracy TS. Comparative modelling of the human UDP-glucuronosyltransferases: insights into structure and mechanism. Xenobiotica. 2007;37: 155–168.

60. Turgeon D, Carrier JS, Lévesque E, Hum DW, Bélanger A. Relative enzymatic activity, protein stability, and tissue distribution of human steroid-metabolizing UGT2B subfamily members. Endocrinology. 2001;142: 778–787.

61. Turgeon D, Carrier J-S, Chouinard S, Bélanger A. Glucuronidation activity of the UGT2B17 enzyme toward xenobiotics. Drug Metab Dispos. 2003;31: 670–676.

62. Zand RS, Jenkins DJ, Diamandis EP. Steroid hormone activity of flavonoids and related compounds. Breast Cancer Res Treat. 2000;62: 35–49.

63. Bushey RT, Dluzen DF, Lazarus P. Importance of UDP-glucuronosyltransferases 2A2 and 2A3 in tobacco carcinogen metabolism. Drug Metab Dispos. 2013;41: 170–179.

64. Gotoh-Saito S, Abe T, Furukawa Y, Oda S, Yokoi T, Finel M, et al. Characterization of human UGT2A3 expression using a prepared specific antibody against UGT2A3. Drug Metab Pharmacokinet. 2019;34: 280–286.

65. Mithöfer A, Boland W. Plant defense against herbivores: chemical aspects. Annu Rev Plant Biol. 2012;63: 431–450.

66. Kuehl GE, Murphy SE. N-glucuronidation of nicotine and cotinine by human liver microsomes and heterologously expressed UDP-glucuronosyltransferases. Drug Metab Dispos. 2003;31: 1361–1368.

67. Mazur A, Lichti CF, Prather PL, Zielinska AK, Bratton SM, Gallus-Zawada A, et al. Characterization of human hepatic and extrahepatic UDP-glucuronosyltransferase enzymes involved in the metabolism of classic cannabinoids. Drug Metab Dispos. 2009;37: 1496– 1504.

68. Li C, Wu Q. Adaptive evolution of multiple-variable exons and structural diversity of drug-metabolizing enzymes. BMC Evol Biol. 2007;7: 69.

69. da Silveira Anacleto TC. Food habits of four armadillo species in the Cerrado area, Mato Grosso, Brazil. ZOOLOGICAL STUDIES-TAIPEI-. 2007;46: 529.

70. Hallström BM, Kullberg M, Nilsson MA, Janke A. Phylogenomic data analyses provide evidence that Xenarthra and Afrotheria are sister groups. Mol Biol Evol. 2007;24: 2059– 2068.

71. Eschler BM, Pass DM, Willis R, Foley WJ. Distribution of foliar formylated phloroglucinol derivatives amongst Eucalyptus species. Biochem Syst Ecol. 2000;28: 813–824.

72. Külheim C, Padovan A, Hefer C, Krause ST, Köllner TG, Myburg AA, et al. The Eucalyptus terpene synthase gene family. BMC Genomics. 2015;16: 450.

