## Supplementary Tables and Figures for "Evolutionary history of mammalian UDP-glucuronosyltransferase (UGT)1 and UGT2 families: the emergence of UGT2B subfamily in eutherians after the diversification of flowering plants"

**This PDF file includes:**

Figures S1 to S8
Tables S1 to S2
Legends for Data S1 to S13

**Other supplementary materials for this manuscript include the following:**

Data S1 to S9

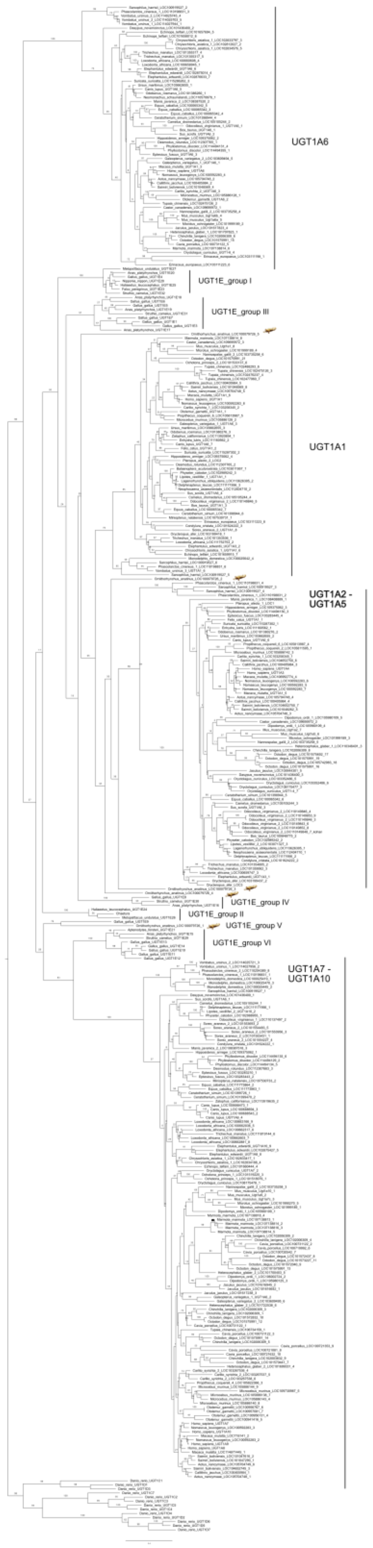


**Fig. S1.** Phylogenetic tree of UDP-glucuronosyltransferase 1 (UGT1) family genes constructed from the amino acid sequence alignments of the 1st exons in mammalian and avian *UGT1* genes. The UGT1E clade names were based on Kawai et al. (2018). Bootstrap values with 100 replicates are shown next to the branches as percentage. The tree is drawn to scale with branch length indicating the expected number of substitutions per site.

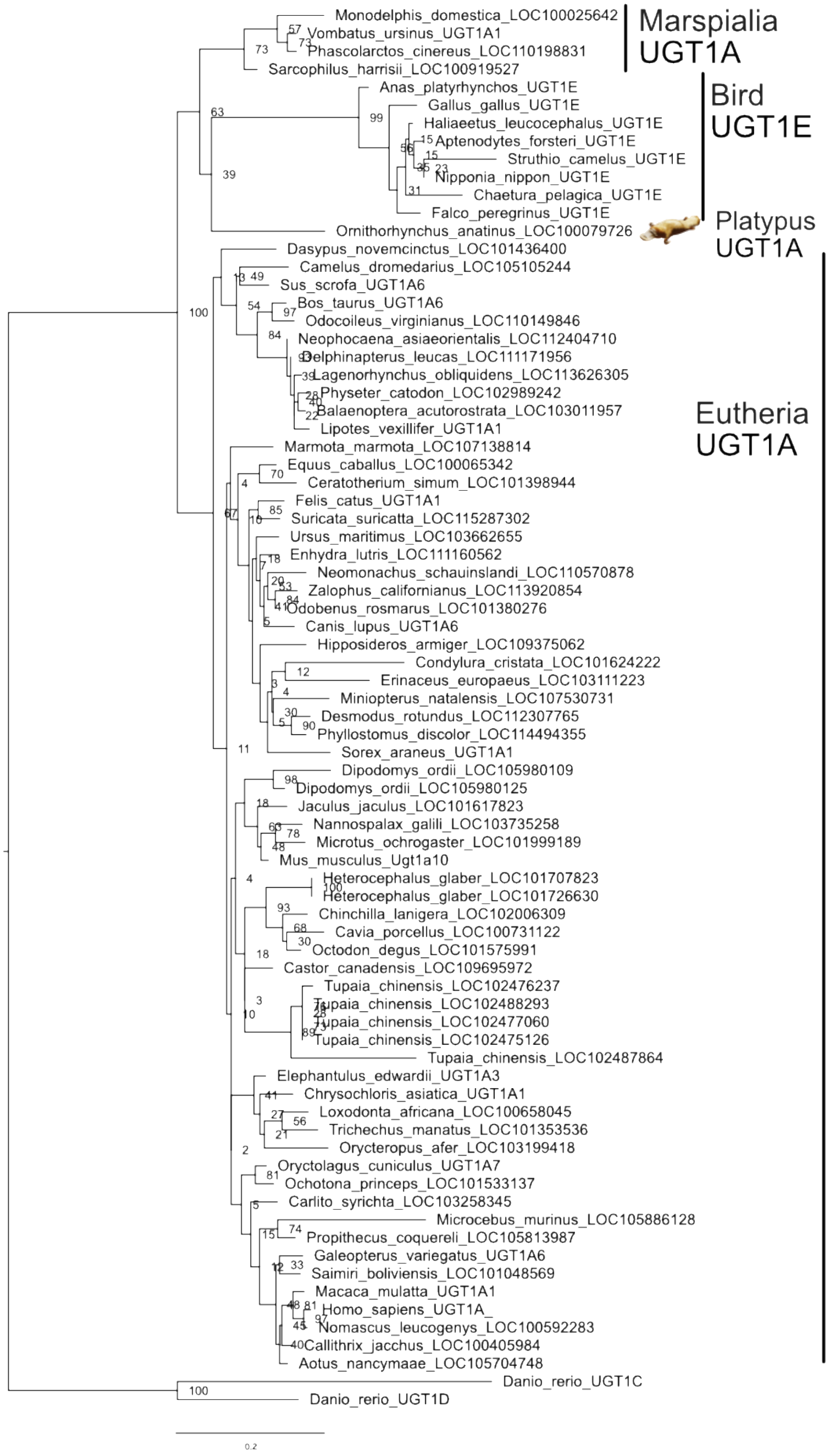


**Fig. S2.** Phylogenetic tree of UDP-glucuronosyltransferase 1 (UGT1) family genes constructed from the amino acid sequence alignments of the 2nd–5th exons in mammalian and avian *UGT*1 genes. Bootstrap values with 100 replicates are shown next to the branches as percentage. The tree is drawn to scale with branch length indicating the expected number of substitutions per site.

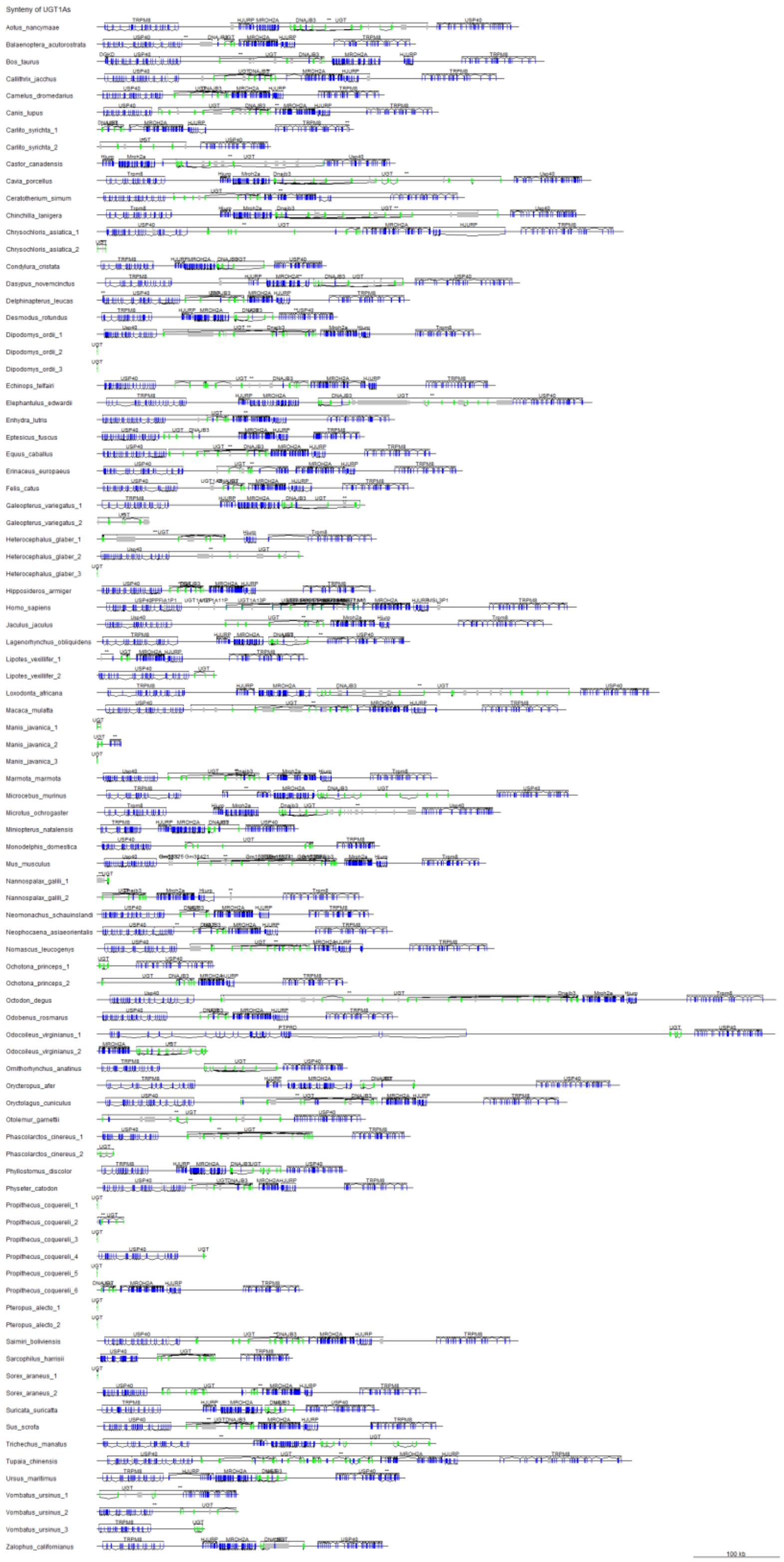


**Fig. S3.** The locus information of the *UGT1A* genes. Blue boxes represent the exons of genes other than *UGT*, green boxes represent the exons of *UGT1A*, and gray boxes represent pseudogenes.

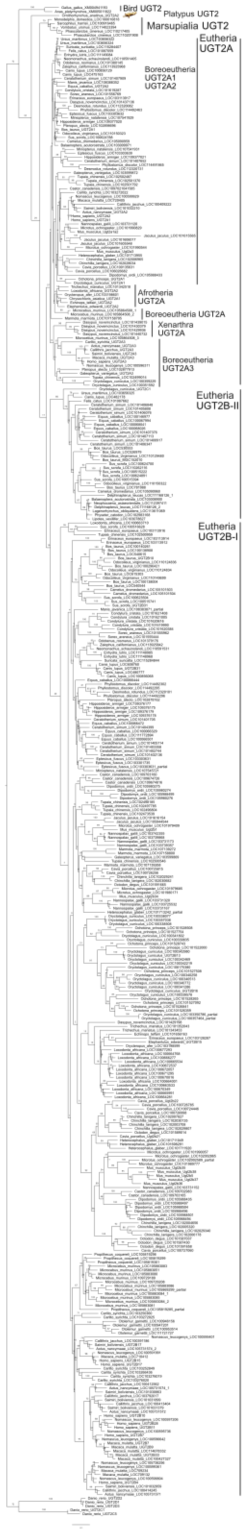


**Fig. S4.** Phylogenetic tree of UDP-glucuronosyltransferase 2 (UGT2) family genes constructed from the amino acid sequence alignments of the 2nd–6th exons in mammalian and avian *UGT2* genes. Bootstrap values with 100 replicates are shown next to the branches as percentage. The tree is drawn to scale with branch length indicating the expected number of substitutions per site.

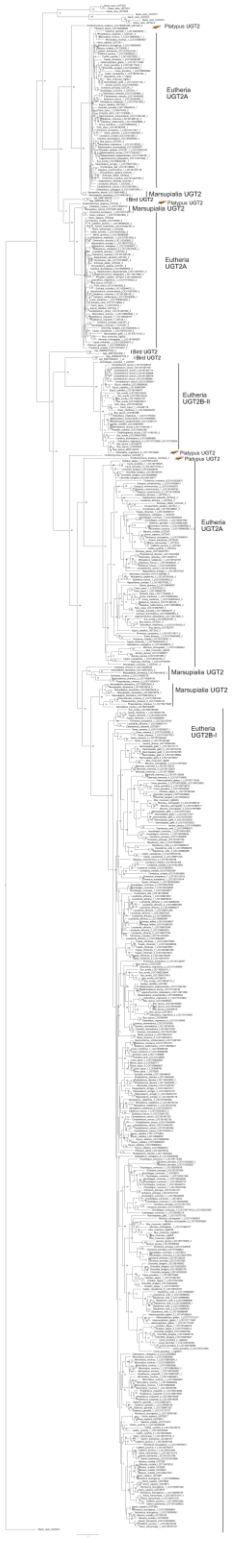


**Fig. S5.** Phylogenetic tree of UDP-glucuronosyltransferase 2 (UGT2) family genes constructed from the amino acid sequence alignments of the 1st exons in mammalian and avian *UGT2* genes. Bootstrap values with 100 replicates are shown next to the branches as percentage. The tree is drawn to scale with branch length indicating the expected number of substitutions per site.

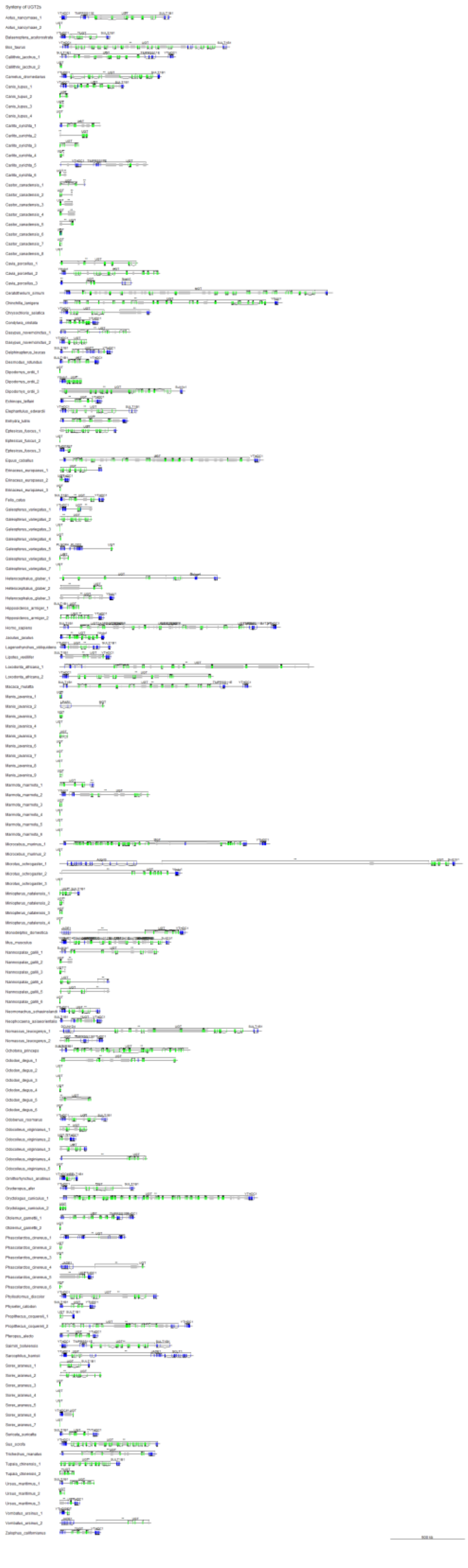

**Fig. S6.** The locus information of the *UGT2* genes. Blue boxes represent the exons of genes other than *UGT*, green boxes represent the exons of *UGT2*, and gray boxes represent pseudogenes.

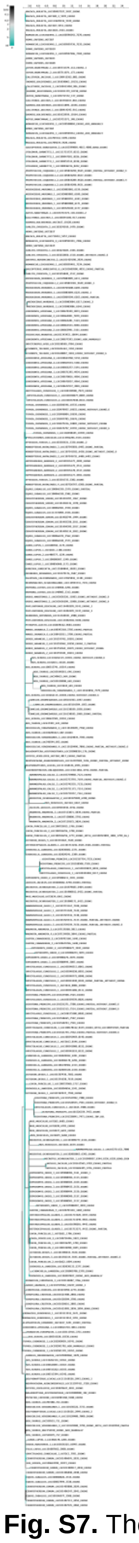

**Fig. S7.** The results of aBSREL analysis in HyPhy for detecting positive selection on UGT2B family genes. The branches with green color show higher omega value and suggest positive selection.

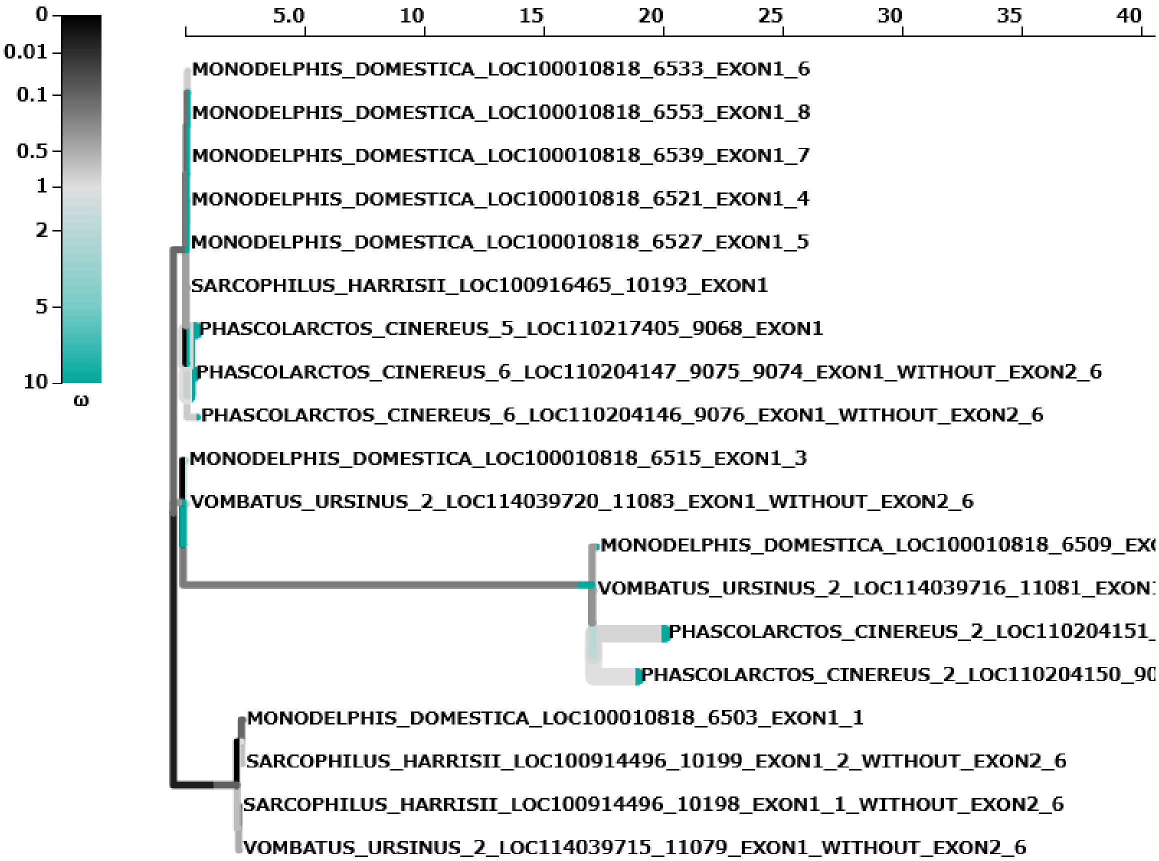


**Fig. S8.** The results of aBSREL analysis in HyPhy for detecting positive selection on *UGT2B* family genes. The branches with green color show higher omega value and suggest positive selection.

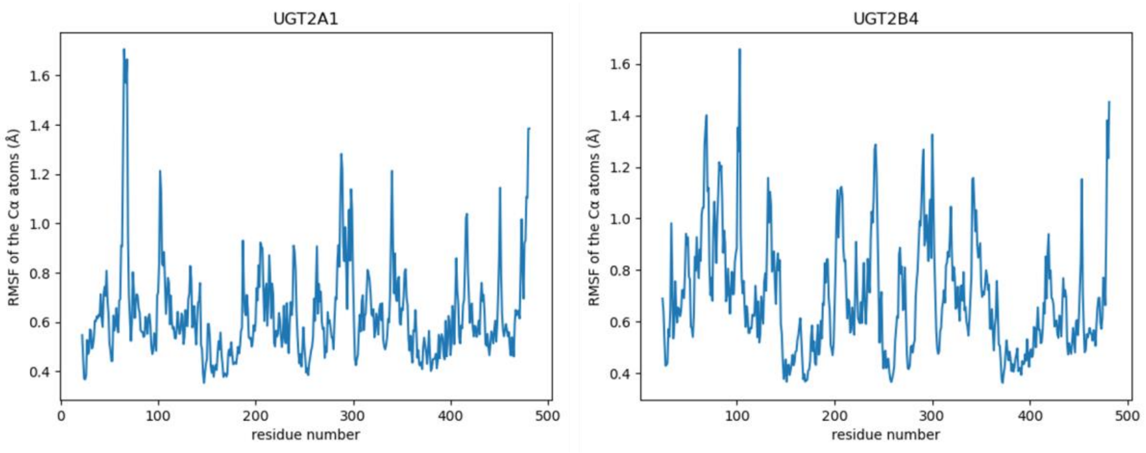


**Fig. S9.** A comparison of RMSF for UGT2A1 and UGT2B4. The graph on the left is for UGT2A1 and one in the right is for UGT2B4. The horizontal axis represents the site of amino acid residues corresponding to UGT2A1 (UniProt accession: P0DTE4) and UGT2B4 (UniProt accession: P06133), while the vertical axis represents the RMSF (Å) of the Cα atoms of the protein backbone.

**Table S1.** Mammalian 67 species used in this study and their feeding habits.

| Scientific_name | Common_name | Family | Feeding habits | Scaffold N50 |
|---|---|---|---|---|
| Aotus_nancymaae | Nancy_Ma's_night_monkey | Aotidae | Omnivorous | 8,268,663 |
| Balaenoptera_acutorostrata | minke_whale | Balaenopteridae | Carnivorous | 12,843,668 |
| Bos_taurus | Cattle | Bovidae | Herbivorous | 103308737 |
| Callithrix_jacchus | Common_marmoset | Callitrichidae | Omnivorous | 5167444 |
| Camelus_dromedarius | Dromedary | Camelidae | Herbivorous | 70369702 |
| Canis_lupus | Dog | Canidae | Carnivorous | 45876610 |
| Carlito_syrichta | Philippine_tarsier | Tarsiidae | Carnivorous | 401181 |
| Castor_canadensis | American_beaver | Castoridae | Herbivorous | 317708 |
| Cavia_porcellus | Guinea_pig | Caviidae | Herbivorous | 7942054 |
| Ceratotherium_simum | White_rhinoceros | Rhinocerotidae | Herbivorous | 26277727 |
| Chinchilla_lanigera | Long-tailed_chinchilla | Chinchillidae | Herbivorous | 21893125 |
| Chrysochloris_asiatica | Cape_golden_mole | Chrysochloridae | Carnivorous | 13470186 |
| Condylura_cristata | Star-nosed_mole | Talpidae | Carnivorous | 55520359 |
| Dasypus_novemcinctus | Nine-banded_armadillo | Dasypodidae | Carnivorous | 1687935 |
| Delphinapterus_leucas | Beluga | Monodontidae | Carnivorous | 31183418 |
| Desmodus_rotundus | Common_vampire_bat | Desmodontidae | Carnivorous | 26869735 |
| Dipodomys_ordii | Ord's_kangaroo_rat | Heteromyidae | Omnivorous | 11931245 |
| Echinops_telfairi | Lesser_hedgehog_tenrec | Tenrecidae | Omnivorous | 54422506 |
| Elephantulus_edwardii | Cape_elephant_shrew | Macroscelididae | Carnivorous | 15011382 |
| Enhydra_lutris | Sea_otter | Mustelidae | Carnivorous | |
| Eptesicus_fuscus | Big_brown_bat | Vespertilionidae | Carnivorous | 13454942 |
| Equus_caballus | Horse | Equidae | Herbivorous | 87230776 |
| Erinaceus_europaeus | Western_European_hedgehog | Erinaceidae | Carnivorous | 3264618 |
| Felis_catus | Cat | Felidae | Carnivorous | 83967707 |
| Galeopterus_variegatus | Sunda_Flying_Lemur | Cynocephalidae | Herbivorous | 245189 |
| Heterocephalus_glaber | Naked_mole_rat | Bathyergidae | Herbivorous | 20532749 |
| Hipposideros_armiger | Great_roundleaf_bat | Hipposideridae | Carnivorous | 2328177 |
| Homo_sapiens | Human | Hominidae | Omnivorous | 67794873 |
| Jaculus_jaculus | Lesser_Egyptian_jerboa | Dipodidae | Herbivorous | 22080993 |
| Lagenorhynchus_obliquidens | Pacific_White-sided_Dolphin | Delphinidae | Carnivorous | 28371583 |
| Lipotes_vexillifer | Yangtze_River_dolphin | Lipotidae | Carnivorous | 2419148 |
| Loxodonta_africana | African_elephant | Elephantidae | Herbivorous | 46401353 |
| Macaca_mulatta | Rhesus_macaque | Cercopithecidae | Omnivorous | 82346004 |
| Manis_javanica | Sunda_pangolin | Manidae | Carnivorous | 204728 |
| Marmota_marmota | Alpine_marmot | Sciuridae | Herbivorous | 31340621 |
| Microcebus_murinus | Gray_mouse_lemur | Cheirogaleidae | Omnivorous | 108171978 |
| Microtus_ochrogaster | Prairie_vole | Cricetidae | Herbivorous | 17270019 |
| Miniopterus_natalensis | Natal_long-fingered_bat | Miniopteridae | Carnivorous | 4315193 |
| Monodelphis_domestica | Gray_short-tailed_opossum | Didelphidae | Omnivorous | 59809810 |
| Mus_musculus | House_mouse | Muridae | Omnivorous | 54517951 |
| Nannospalax_galili | Upper_Galilee_mountains_blind_mole_rat | Spalacidae | Herbivorous | 3618479 |
| Neomonachus_schauinslandi | Hawaiian_monk_seal | Phocidae | Carnivorous | 29518589 |
| Neophocaena_asiaeorientalis | Narrow-ridged_finless_porpoise | Phocoenidae | Carnivorous | 6341296 |
| Nomascus_leucogenys | Northern_white-cheeked_gibbon | Hylobatidae | Omnivorous | 35886894 |
| Ochotona_princeps | American_pika | Ochotonidae | Herbivorous | 26863993 |
| Octodon_degus | Common_degu | Octodontidae | Herbivorous | 12091372 |
| Odobenus_rosmarus | Walrus | Odobenidae | Carnivorous | 2616778 |
| Odocoileus_virginianus | White-tailed_deer | Cervidae | Herbivorous | |
| Ornithorhynchus_anatinus | Platypus | Ornithorhynchidae | Carnivorous | 83338043 |
| Orycteropus_afer | Aardvark | Orycteropodidae | Carnivorous | |
| Oryctolagus_cuniculus | European_rabbit | Leporidae | Herbivorous | 35972871 |
| Otolemur_garnettii | Northern_greater_galago | Galagidae | Omnivorous | 13852661 |
| Phascolarctos_cinereus | Koala | Phascolarctidae | Herbivorous | |
| Phyllostomus_discolor | Pale_spear-nosed_bat | Phyllostomidae | Omnivorous | 110241909 |
| Physeter_catodon | Sperm_Whale | Physeteridae | Carnivorous | 122182240 |
| Propithecus_coquereli | Coquerel's_sifaka | Indriidae | Herbivorous | 5604909 |
| Pteropus_alecto | Black_flying_fox | Pteropodidae | Herbivorous | 15954802 |
| Saimiri_boliviensis | Black-capped_Squirrel_Monkey | Cebidae | Omnivorous | 18744880 |
| Sarcophilus_harrisii | Tasmanian_devil | Dasyuridae | Carnivorous | 611347268 |
| Sorex_araneus | Common_shrew | Soricidae | Carnivorous | 22794405 |
| Suricata_suricatta | Meerkat | Herpestidae | Omnivorous | 141453419 |
| Sus_scrofa | Boar | Suidae | Omnivorous | 88231837 |
| Trichechus_manatus | Caribbean_manatee | Trichechidae | Herbivorous | 14442683 |
| Tupaia_chinensis | Chinese_tree_shrew | Tupaiidae | Omnivorous | 3670124 |
| Ursus_maritimus | Polar_bear | Ursidae | Carnivorous | 15940661 |
| Vombatus_ursinus | Common_wombat | Vombatidae | Herbivorous | 28503419 |
| Zalophus_californianus | California_sea_lion | Otariidae | Carnivorous | 143424588 |

**Table S2.** Models for maximum likelihood phylogenetic analysis.

| Subject | Model |
|---|---|
| UGT1A 1st exons | JTT+F_Gamma |
| UGT1A 2nd-5th exons | LG4X_Gamma |
| UGT2 1st exons | HIVb+F_Gamma |
| UGT2 2nd-6th exons | HIVb_Gamma |

**Table S3.** Predicted protein structure described in this study and ERRAT value. Protein ID shows

| GenBank accession | UGT subfamily | species | ERRAT value | desctribed structure in this study |
|---|---|---|---|---|
| XP_004471524.1 | UGT1A | armadillo | 84.2912 | helix |
| XP_012385649.1 | UGT1A | armadillo | 86.0153 | helix |
| XP_012385650.1 | UGT1A | armadillo | 84.9609 | helix |
| XP_004477619.1 | UGT2A | armadillo | 91.3215 | helix |
| XP_012374729.1 | UGT2A | armadillo | 85.9345 | helix |
| XP_004477626.1 | UGT2A | armadillo | 90.9944 | hinge |
| XP_004478400.1 | UGT2A | armadillo | 92.3364 | hinge |
| XP_004478401.1 | UGT2A | armadillo | 90.3475 | helix |
| XP_004478398.1 | UGT2B | armadillo | 91.9386 | hinge |
| XP_023413506.1 | UGT1A | elephant | 83.5766 | helix |
| XP_023413508.1 | UGT1A | elephant | 86.0075 | helix |
| XP_023413502.1 | UGT1A | elephant | 84.749 | helix |
| XP_023413503.1 | UGT1A | elephant | 84.6154 | helix |
| XP_010595963.2 | UGT1A | elephant | 84.913 | helix |
| XP_023413460.1 | UGT1A | elephant | 85.206 | helix |
| XP_003418002.1 | UGT1A | elephant | 87.3346 | helix |
| XP_003418003.1 | UGT1A | elephant | 84.7458 | helix |
| XP_023413461.1 | UGT1A | elephant | 88.6827 | helix |
| XP_010592459.1 | UGT2A | elephant | 86.8979 | hinge |
| XP_010592460.2 | UGT2A | elephant | 90.4398 | hinge |
| XP_003414218.1 | UGT2A | elephant | 91.619 | helix |
| XP_003414217.1 | UGT2A | elephant | 87.2047 | helix |
| XP_023409136.1 | UGT2B | elephant | 91.1368 | hinge |
| XP_010592496.1 | UGT2B | elephant | 92.5806 | seems not functional |
| XP_003414219.1 | UGT2B | elephant | 93.3981 | hinge |
| XP_010592468.2 | UGT2B | elephant | 90.7721 | hinge |
| XP_010592474.2 | UGT2B | elephant | 86.6795 | hinge |
| XP_003414225.1 | UGT2B | elephant | 93.0368 | hinge |
| XP_010592470.2 | UGT2B | elephant | 90.1734 | hinge |
| XP_023410886.1 | UGT2B | elephant | 91.6828 | hinge |
| XP_023410882.1 | UGT2B | elephant | 91.7148 | hinge |
| XP_023410885.1 | UGT2B | elephant | 88.0539 | hinge |
| XP_003415960.1 | UGT2B | elephant | 90.1544 | hinge |

| | | | | |
|---|---|---|---|---|
| XP_023410881.1 | UGT2B | elephant | 93.0097 | hinge |
| XP_010594014.1 | UGT2B | elephant | 93.8104 | hinge |
| XP_023410871.1 | UGT2B | elephant | 91.8447 | hinge |
| XP_023410868.1 | UGT2B | elephant | 90.1734 | hinge |
| XP_016284843.1 | UGT1A | opossum | 89.5551 | helix |
| XP_007485971.1 | UGT1A | opossum | 90.3846 | helix |
| XP_001376379.2 | UGT1A | opossum | 88.9306 | helix |
| XP_001376361.2 | UGT1A | opossum | 84.8077 | helix |
| XP_007496418.1 | UGT2A | opossum | 85.9649 | helix |
| XP_007496419.2 | UGT2B | opossum | 85.6031 | helix |
| XP_007496420.1 | UGT2B | opossum | 91.8919 | hinge |
| XP_007496422.1 | UGT2B | opossum | 88.8672 | hinge |
| XP_007496423.1 | UGT2B | opossum | 89.5551 | hinge |
| XP_007496424.2 | UGT2B | opossum | 84.9206 | hinge |
| XP_007496425.1 | UGT2B | opossum | 89.4325 | helix |
| XP_016278706.1 | UGT2B | opossum | 88.3495 | hinge |
| XP_007666942.1 | UGT1A | platypus | 90.9789 | helix |
| XP_001510659.3 | UGT1A | platypus | 87.5486 | helix |
| XP_007666949.2 | UGT1A | platypus | 88.2466 | helix |
| XP_007666935.1 | UGT1A | platypus | 88.7405 | helix |
| XP_028924820.1 | UGT1A | platypus | 85.6597 | helix |
| XP_028929290.1 | UGT2 | platypus | 91.0985 | hinge |
| XP_028929292.1 | UGT2 | platypus | 94.084 | helix |
| XP_028929294.1 | UGT2 | platypus | 89.8833 | helix |
| XP_028929529.1 | UGT2 | platypus | 88.7405 | helix |

**Dataset S1 (separate file).** Sequences used as queries.

**Dataset S2 (separate file).** Sequences used for phylogenetic analysis, including the 1st exon of UGT1A, the 2nd-5th exon of UGT1A, the 1st exon of UGT2, and the 2nd-6th exon of UGT2. The alignments are separated by ### respectively.

**Dataset S3 (separate file).** The url links to entrez gene of the gene used in the analysis and whether it is a pseudogene or not.

**Dataset S4 (separate file).** Sequence and phylogenetic tree of *UGT1A6* used for RELAX analysis in HyPhy.

**Dataset S5 (separate file).** Sequence of marsupial *UGT2* family genes used for aBSREL analysis in HyPhy.

**Dataset S6 (separate file).** Sequence of eutherian *UGT2A* family genes used for aBSREL analysis in HyPhy.

**Dataset S7 (separate file).** Sequence and phylogenetic tree of marsupial *UGT2B* family genes used for aBSREL analysis in HyPhy.

**Dataset S8 (separate file).** Sequences of mammalian *UGT1A* genes used for FUBAR analysis in Hyphy.

**Dataset S9 (separate file).** Sequences of eutherian *UGT2B* genes used for FUBAR analysis in Hyphy.

**Dataset S10 (separate file).** Structure information of human UGT2A1 and MD simulation parameters used for OpenMM.

**Dataset S11 (separate file).** Structure information of human UGT2B4 and MD simulation parameters used for OpenMM.

**Dataset S12 (separate file).** The results of aBSREL analysis in HyPhy for detecting positive selection on *UGT2B* family genes.

**Dataset S13 (separate file).** The results of aBSREL analysis in HyPhy for detecting positive selection on Marsupialia *UGT2* family genes.
